# Untangling dopamine and glutamate in the ventral tegmental area

**DOI:** 10.1101/2025.02.25.640201

**Authors:** Emily D. Prévost, Lucy A. Ward, Daniel Alas, Giulia Aimale, Sara Ikenberry, Katie Fox, Julianne Pelletier, Annie Ly, Jayson Ball, Zachary P. Kilpatrick, Kailyn Price, Abigail M. Polter, David H. Root

## Abstract

Ventral tegmental area (VTA) dopamine neurons are of great interest for their central roles in motivation, learning, and psychiatric disorders. While hypotheses of VTA dopamine neuron function posit a homogenous role in behavior (e.g., prediction error), they do not account for molecular heterogeneity. We find that glutamate-dopamine, nonglutamate-dopamine, and glutamate-only neurons are dissociable in their signaling of reward and aversion-related stimuli, prediction error, and electrical properties. In addition, glutamate-dopamine and nonglutamate-dopamine neurons differ in dopamine release dynamics. Aversion-related recordings of all dopamine neurons (not considering glutamate co-transmission) showed a mixed response that obscured dopamine subpopulation function. Within glutamate-dopamine neurons, glutamate and dopamine release had dissociable contributions toward reward and aversion-based learning and performance. Based on our results, we propose a new hypothesis on VTA dopamine neuron function: that dopamine neuron signaling patterns and their roles in motivated behavior depend on whether or not they co-transmit dopamine with glutamate.

## INTRODUCTION

Dopaminergic neurons in the ventral tegmental area (VTA) regulate motivation-related behaviors as well as contribute to psychiatric disorders via their influence on attaining rewards or avoiding punishment^1^. Though many different ideas have been proposed to explain the functions of VTA dopamine neurons, such as prediction error^2^, causal learning^3^, associative learning^4,5^, and others^6-12^, they are all based on the premise of a homogeneous population of dopamine neurons. However, it is well known that VTA neurons are molecularly diverse^1,13^, and it is unclear how molecular heterogeneity contributes to the wide range of functions ascribed to VTA dopamine neurons. A primary molecular distinction between subtypes of VTA dopamine neurons is whether they co-transmit dopamine and glutamate or release dopamine without glutamate^14,15^. Yet, the roles of glutamate-dopamine and nonglutamate-dopamine neurons in motivated behavior are unknown. Moreover, the individual contributions of glutamate and dopamine release by glutamate-dopamine co-transmitting neurons on learning to attain rewards or avoid aversive stimuli are unknown. Further, because VTA glutamate neurons can co-transmit glutamate with dopamine or co-transmit glutamate with GABA, it has been challenging to untangle glutamate and dopamine function of VTA neurons, especially the unknown functions of VTA neurons that release *only* glutamate. Here, we parse glutamate-dopamine neurons from nonglutamate-dopamine neurons and glutamate-only neurons, as well as identify the roles of glutamate and dopamine release from glutamate-dopamine co-transmitting neurons toward Pavlovian-guided reward- and aversion-related motivated behaviors. Our strategy involved combining intersectional and subtractive genetics with *ex vivo* whole-cell recordings, *in vivo* calcium recordings, optogenetics, *in vivo* dopamine sensors, and cell-type and neurotransmitter-specific knockdown. We find that glutamate-dopamine, nonglutamate-dopamine, and glutamate-only neurons are dissociable in their signaling of cued rewards, cued and uncued aversive stimuli, and reward prediction error, and have separable electrical properties and excitability. In addition, glutamate-dopamine and nonglutamate-dopamine neurons differ in their dopamine release dynamics in their preferential projections to nucleus accumbens medial shell and core, respectively. Experiments recording all dopamine neurons without considering glutamate co-transmission showed a signaling pattern in response to aversive stimuli that was the approximate average of glutamate-dopamine and nonglutamate-dopamine neuronal activities, suggesting separation of glutamate-dopamine and nonglutamate-dopamine neurons is an important factor for identifying different aversion-related VTA dopamine functions. Examining glutamate-dopamine neurons further, we found that while glutamate release from glutamate-dopamine neurons was necessary for the vigor of attaining rewards without affecting learning, dopamine release was necessary for learning about stimuli related to negative or aversive events without effects on attaining rewards. We also present a neural circuit model incorporating dopaminergic subtypes which recapitulates the roles of distinct neurotransmitters from glutamate-dopamine co-transmitting neurons. Based on our results, we propose a new hypothesis of VTA dopamine neuron function: that dopamine neuron signaling patterns and roles in motivated behavior depend on whether they co-transmit dopamine with glutamate or release dopamine without glutamate.

## RESULTS

### Intersectional and subtractive cell-type targeting strategies

To gain genetic access to VTA glutamate-dopamine, nonglutamate-dopamine, and glutamate-only neurons, we generated two double transgenic mouse lines. For dopamine subtypes, VGluT2::Cre mice were crossed with TH::Flp mice. Double transgenic VGluT2::Cre/TH::Flp offspring were injected in VTA with INTRSECT AAVs that expressed GCaMP6m in neurons expressing both Cre and Flp (Con/Fon-GCaMP) to label glutamate-dopamine neurons, or in neurons expressing Flp without Cre (Coff/Fon-GCaMP) to label nonglutamate-dopamine neurons (**Figure 1A**). To compare dopamine subpopulation results to all dopamine neurons, as is the standard in the field, a separate group of single recombinase TH::Flp mice were injected in VTA with Coff/Fon-GCaMP to label all dopamine neurons without taking into account whether these neurons release dopamine alone or together with glutamate (**Figure 1A**). To gain genetic access to VTA glutamate-only neurons, we used a strategy to label neurons that explicitly do not release GABA or dopamine, leaving the remaining neurons as those that release glutamate without co-releasing dopamine or GABA. To accomplish this, we crossed VGaT::Cre mice with TH::Flp mice. In double transgenic VGaT::Cre/TH::Flp offspring, we injected in VTA an INTRSECT AAV that expressed GCaMP6m in neurons lacking either Cre or Flp: (Cre-Or-Flp)Off-GCaMP6m (COFO-GCaMP6m) (**Figure 1A**). Consistent with prior localizations of VGluT2 and TH expressing VTA neurons^15-21^, nonglutamate-dopamine neurons were found laterally (**Figure 1B-F**), glutamate-dopamine neurons were near the midline, and both nonglutamate-dopamine and glutamate-dopamine neurons comingled at the boundary of medial and lateral VTA (**Figure 1G-F**). We profiled the GCaMP-labeled neurons for the expression of transcripts encoding VGluT2 and TH. In VGluT2::Cre/TH::Flp mice injected in VTA with Con/ Fon vectors targeting glutamate-dopamine neurons, 96.02 ± 1.9% of GCaMP-labeled neurons expressed both VGluT2 and TH mRNA (1142/1207 neurons from 3 mice) and very few neurons (3.42 ± 1.7%) expressed TH without VGluT2 mRNA (**Figure 1L**). In contrast, for VGluT2::Cre/TH::Flp mice injected in VTA with Coff/ Fon vectors targeting nonglutamate-dopamine neurons, 97.4 ± 0.72% of GCaMP-labeled neurons expressed TH without VGluT2 mRNA (671/690 neurons from 3 mice) and very few neurons (2.49 ± 0.69%) expressed both VGluT2 and TH mRNA (**Figure 1L**). For VGaT::Cre/ TH::Flp mice injected in VTA with COFO-GCaMP6m to target glutamate-only neurons, labeled neurons were most often observed around the midline, consistent with prior literature^15,17,19,22^ (**Figure 1M-F**). We profiled the GCaMP-labeled neurons for the expression of transcripts encoding VGluT2 and VGaT, as well as TH immunoreactivity (TH-IR). About 90% of GCaMP-labeled neurons expressed VGluT2 mRNA without VGaT mRNA or TH-IR co-expression (89.9 ± 0.59%, 505/563 neurons from 4 mice) (**Figure 1S**). Other cell-types labeled were sparse, varying between 0-3.15% of GCaMP-labeled neurons, which likely reflects a low population of Cre- or Flp-lacking VTA neurons in the VGaT::Cre/TH::Flp line. Together, INTRSECT vectors effectively target the proposed dopaminergic and glutamatergic cell-types.

**Figure 1.**
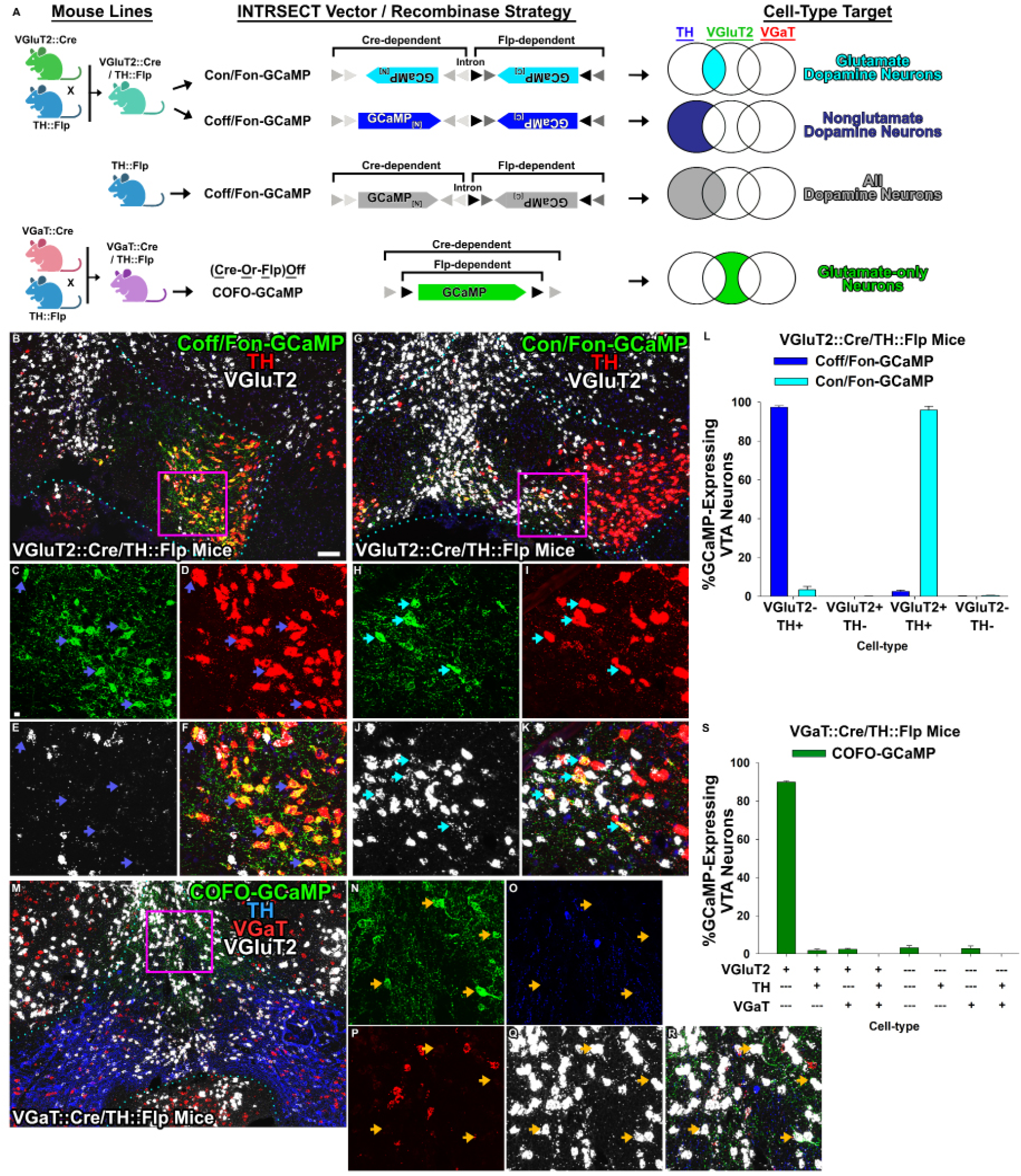
Genetic access to VTA dopaminergic and glutamatergic cell-types. **A**. Neuronal targeting strategies. VGluT2::Cre mice were crossed with TH::Flp mice. VGluT2::Cre/TH::Flp offspring were injected in VTA with INTRSECT AAVs encoding GCaMP6m in a Cre AND Flp manner (Con/Fon) to target glutamate-dopamine neurons or a NOT Cre AND Flp manner (Coff/Fon) to target nonglutamate-dopamine neurons. In TH::Flp mice, Coff/Fon vectors target all dopamine neurons (glutamate-dopamine neurons and nonglutamate-dopamine neurons). To target glutamate-only neurons, VGaT::Cre mice were crossed with TH::Flp mice. VGaT::Cre/ TH::Flp offspring were injected in VTA with an INTRSECT AAV encoding GCaMP6m in a NOT(Cre or Flp) manner to select neurons that do not singularly release or co-release GABA or dopamine, which in VTA selects neurons that only release glutamate. C = Cre, F = Flp, on = must express, off = must not express. [N] = N terminus, [C] = C terminus. Gray triangles = loxP (light) and lox2722 (dark). Black triangles = F3 (black) and F5 (dark). **B-F**. Coff/Fon GCaMP neurons were examined for TH mRNA (red) and VGluT2 mRNA (white). Blue arrows are example GCaMP TH+ VGluT2-neurons. **G-K**. Con/Fon-GCaMP6m neurons were examined for TH mRNA (red) and VGluT2 mRNA (white). Cyan arrows are example GCaMP TH+ VGluT2+ neurons. **L**. Quantification of GCaMP expression. **M-R**. VGaT::Cre/TH::Flp mice were injected in VTA with COFO-GCaMP6m. GCaMP-labeled neurons were examined for TH protein (blue), VGaT mRNA (red), and VGluT2 mRNA (white). Orange arrows are example GCaMP VGluT2+ TH-VGaT-neurons. **S**. Quantification of GCaMP expression. Scale bar in B = 100 µm, applies to G, M. Scale bar in C = 10 µm, applies to C-F, H-K, N-R.

### Cell-type specific ex vivo electrophysiological pro**perties**

We first examined whether VTA glutamate-dopamine, nonglutamate-dopamine, and glutamate-only neurons exhibited cell-type specific electrophysiological properties *ex vivo* (**Figure 2**). VGluT2::Cre/TH::Flp mice were injected in VTA with a mixture of AAVs encoding Con/Fon-eYFP and Coff/ Fon-mCherry to label glutamate-dopamine neurons in eYFP and nonglutamate-dopamine neurons in mCherry. VGaT::Cre/TH::Flp mice were injected in VTA with an AAV encoding COFO-oScarlet to label glutamate-only neurons in oScarlet. Fluorescently labeled neurons were then recorded *ex vivo*. Historically, features such as the presence of an I_h_, high capacitance, presence of dopamine D2 receptors, and pacemaker firing with broad action potentials have been used as electrophysiological markers of dopaminergic neurons, although the reliability of these measures has been called into question^23,24^. We therefore examined distribution of these and other electrophysiological properties. Capacitance and input resistance did not differ between the three groups, while the resting membrane potential was more hyperpolarized in glutamate-only neurons relative to the other two groups (**Supplementary Table 1**). I_h_ was most frequently observed in glutamate-only neurons (61.5% of cells), as well as in a number of nonglutamate-dopamine neurons (35.7% of cells) (**Figure 2A-F**). However, detectable I_h_ was found in very few glutamate-dopamine cells (11.1%). Whole-cell spontaneous action potentials also differed between the three cell-types (**Figure 2D-F**). Action potential half-width was broadest in nonglutamate-dopamine neurons and narrowest in glutamate-only neurons, while glutamate-dopamine neurons showed intermediate half-widths (**Figure 2D-F**). The magnitude of the after-hyperpolarization was significantly smaller in glutamate-dopamine neurons than in the other two cell-types (**Figure 2F**). Spontaneous cell-attached firing rates were highest in glutamate-only neurons and lowest in nonglutamate-dopamine neurons, with firing rates of glutamate-dopamine neurons again intermediate between the two (**Figure 2G-F**). While cell-attached firing of at least some cells from each cell-type was inhibited in response to bath application of the dopamine D2 receptor agonist quinpirole (1 µM), D2 receptor sensitivity was universal among nonglutamate-dopamine neurons but only found in 55-60% of glutamate-dopamine and glutamate-only neurons (**Figure 2I**). Glutamate-only and glutamate-dopamine neurons showed greater excitability than nonglutamate-dopamine neurons in response to increasing depolarizing current injections (**Figure 2J, L**). Finally, the majority of glutamate-only neurons (78%) showed rebound firing of action potentials in response to hyperpolarization, while very few (8%) or no cells exhibited this property in glutamate-dopamine or nonglutamate-dopamine neurons, respectively. Altogether, VTA glutamate-dopamine, nonglutamate-dopamine, and glutamate-only subpopulations exhibit holistically independent constellations of electrophysiological characteristics.

**Figure 2.**
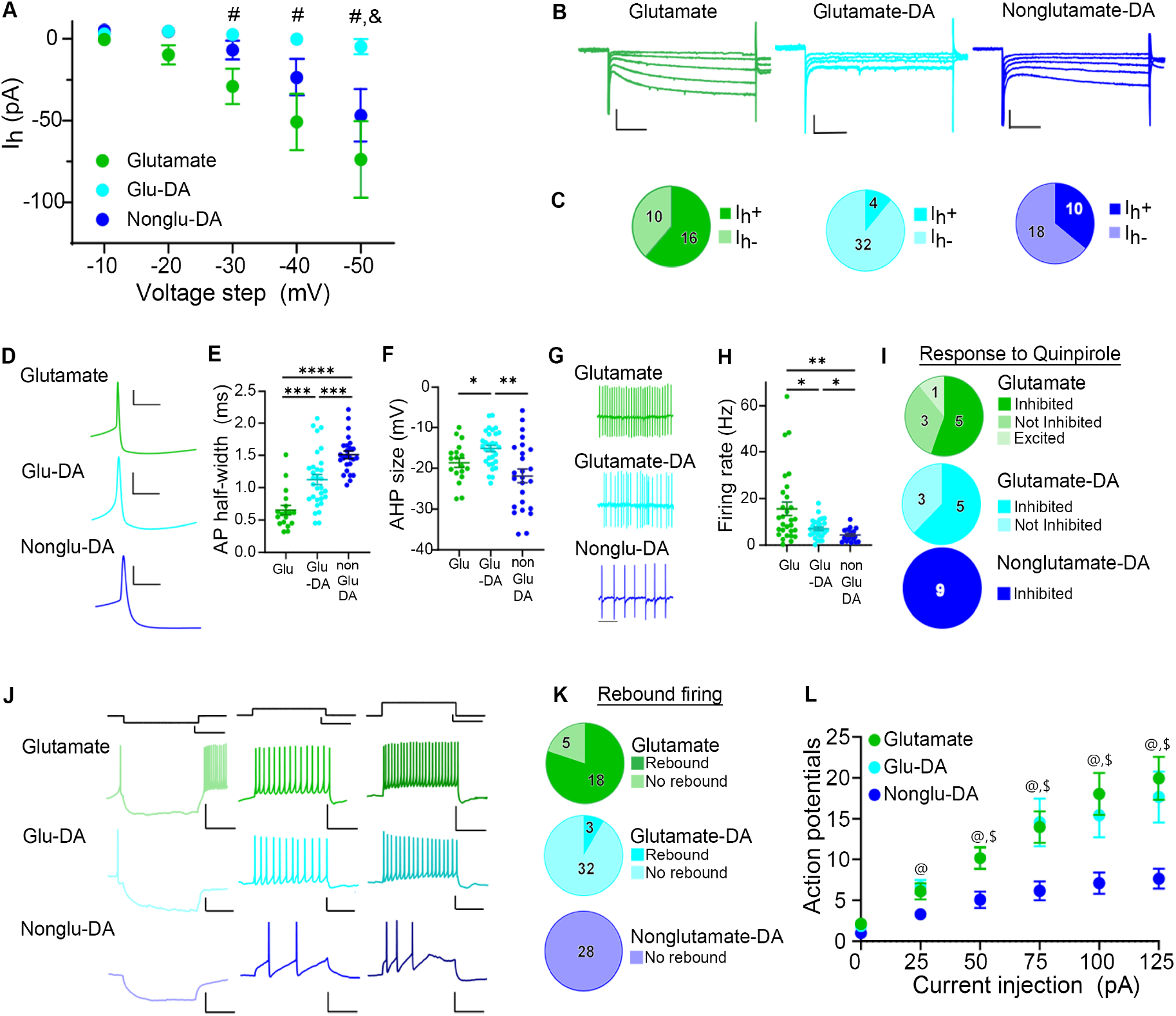
Electrophysiological access to VTA dopaminergic and glutamatergic cell-types. **A**. Hyperpolarization-induced currents (I_h_) across a range of hyperpolarizing steps. Two-way repeated measure ANOVA, cell-type x voltage step *F*(8, 360) = 5.57, *p* < 0.001. # *p* < 0.05, Tukey’s Multiple Comparison Test Glutamate vs. Glu-DA; & *p* < 0.05 Tukey’s Multiple Comparison Test, Non-Glu DA vs Glu-DA. **B**. Representative traces of hyperpolarization-induced currents from each cell-type. Scale bars 100 pA, 100 ms. **C**. Pie charts representing the number of cells that were I_h_+ (maximal current >25 pA) or I_h_- (maximal current <25 pA). Inset numbers represent the number of cells in each category. χ^2^ (17.38, 2), *p* < 0.001. **D**. Representative averages of twenty spontaneous action potentials. Scale bars 20 mV, 10 ms. **E**. Half-width of spontaneous action potentials measured in whole-cell configuration. One-way ANOVA, *F*(2, 71) = 29.34, *p* < 0.0001. *** *p* < 0.001; **** *p* < 0.0001, Tukey’s Multiple Comparisons test. **F**. AHP magnitude of spontaneous action potentials measured in whole-cell configuration. Welch’s ANOVA test *W*(2, 39.18) = 7.98, *p* = 0.001. * *p* < 0.05; ** *p* < 0.01, Dunnett’s multiple comparisons test. **G**. Representative traces of cell-attached action potential firing from each cell-type. Scale bar = 500 ms. **H**. Cell-attached firing rate. Welch’s ANOVA test *W*(2, 47.04) = 10.05, *p* < 0.001. * *p* < 0.05; ** *p* < 0.01, Dunnett’s multiple comparisons test. **I**. Pie charts representing the number of cells where the cell-attached firing rate was inhibited, excited, or unaffected by bath application of 1 uM quinpirole. Inset numbers represent the number of cells in each category. Fisher’s exact test, *p* = 0.100. **J**. Representative traces of evoked action potentials in response to -50 pA, 50 pA, and 100 pA current injections. Scale bars 50 pA, 100 ms (top row); 10 mV, 100 ms (bottom three rows). **K**. Pie charts representing the number of cells that exhibited rebound firing of action potentials following hyperpolarization. Inset numbers represent the number of cells in each category. Fisher’s exact test, *p* < 0.001. **L**. Input-output function of action potentials generated by depolarizing current injections. Mixed-effects model *F*(10, 369) = 4.37, *p* < 0.001. @ *p* < 0.05; Glu-DA vs non-glu DA; $ *p* < 0.05, glutamate vs. non-glu DA.

### Cell-type specific in vivo calcium signaling dynamics in motivated behaviors

To examine whether VTA cell-types differ in motivated behavior-related signaling, we expressed GCaMP6m in either VTA glutamate-dopamine neurons, nonglutamate-dopamine neurons, glutamate-only neurons, or all dopamine neurons and implanted an optic fiber in VTA (**Supplementary Figure 1**). Mice were trained to associate an auditory stimulus (CS+) with sucrose reward delivery and a different auditory stimulus (CS-) with no reward delivery (**Figure 3A**). All groups learned to significantly increase reward port entries during the CS+ (CS+ hits) and decrease reward port entries during the CS-(CS-hits; **Figure 3B-F**). Sidak-adjusted pairwise comparisons showed that while some statistically significant differences were observed between groups during the earliest training days, no group differences in CS+ hits or CS-hits were observed during the last four days of training before neuronal recordings (**Figure 3B**). Following training, GCaMP mice were recorded during the same paradigm with the exception that 11% of trials presented the CS+ cue but reward was omitted to probe for cell-types that signaled reward prediction error.

**Figure 3.**
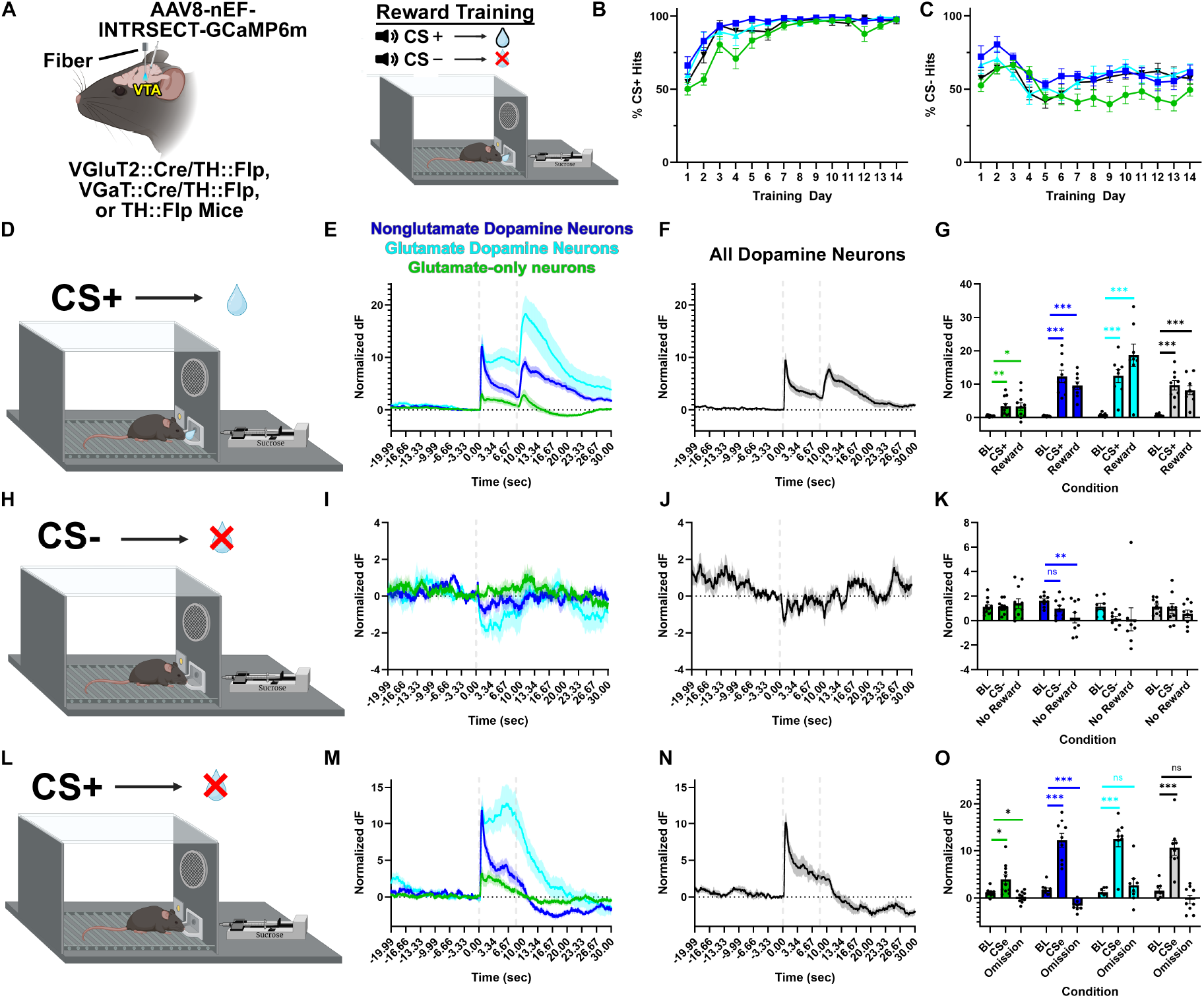
Cell-type specific cued reward-related signaling. **A**. Experimental setup. **B-C**. Percent of reward port entries during CS+ trials (**B**) or CS-trials (**C**) over training days. CS+ hits day x group interaction *F*(39, 429) = 2.06, *p* = 0.020; CS-hits day x group interaction *F*(39, 429) = 1.68, *p* = 0.008. **D-G**. Neuronal activity during CS+ and reward delivery trials where mice entered the reward port. Nonglutamate-dopamine (dark blue), glutamate-dopamine (cyan), glutamate-only (green), and all dopamine (black) neuronal activity. Glutamate-only neurons: main effect of epoch *F*(2,20) = 8.27, *p* = 0.012, posthoc contrast tests: BL x CS+ *F*(1,10) = 13.64, *p* = 0.004; BL x reward *F*(1,10) = 6.81, *p* = 0.026. Nonglutamate-dopamine neurons: main effect of epoch *F*(2,16) = 40.17, *p* < 0.001, posthoc contrast tests: BL x CS+ *F*(1,8) = 41.52, *p* < 0.001; BL x reward *F*(1,8) = 62.11, *p* < 0.001. Glutamate-dopamine neurons: main effect of epoch *F*(2,14) = 22.65, *p* < 0.001, posthoc contrast tests: BL x CS+ *F*(1,7) = 37.67, *p* < 0.001; BL x reward *F*(1,7) = 29.52, *p* < 0.001. All dopamine neurons: main effect of epoch *F*(2,16) = 34.13, *p* < 0.001, posthoc contrast tests: BL x CS+ *F*(1,8) = 42.19, *p* < 0.001; BL x reward *F*(1,8) = 33.45, *p* < 0.001. **H-K**. Neuronal activity during CS- and no reward delivery trials where mice entered the reward port. Glutamate-only neurons: *F*(2,20) = 0.52, *p* = 0.601. Nonglutamate-dopamine neurons: main effect of epoch *F*(2,16) = 7.95, *p* = 0.004, posthoc contrast tests: BL x CS-*F*(1,8) = 4.10, *p* = 0.077; BL x no reward *F*(1,8) = 11.65, *p* = 0.009. Glutamate-dopamine neurons: *F*(2,14) = 0.99, *p* = 0.398. All dopamine neurons: *F*(2,16) = 0.73, *p* = 0.499. **L-O**. Neuronal activity during CSe and reward omission trials where mice entered the reward port. Glutamate-only neurons: *F*(2,20) = 10.77, *p* = 0.005, posthoc contrast tests: BL x CSe *F*(1,10) = 8.92, *p* = 0.014; BL x omission *F*(1,10) = 5.21, *p* = 0.046. Nonglutamate-dopamine neurons: main effect of epoch *F*(2,16) = 60.09, *p* < 0.001, posthoc contrast tests: BL x CSe *F*(1,8) = 57.91, *p* < 0.001; BL x omission *F*(1,8) = 38.54, *p* < 0.001. Glutamate-dopamine neurons: main effect of epoch *F*(2,14) = 28.36, *p* < 0.001, posthoc contrast tests: BL x CSe *F*(1,7) = 45.23, *p* < 0.001; BL x omission *F*(1,7) = 1.12, *p* = 0.325. All dopamine neurons: main effect of epoch *F*(2,16) = 31.12, *p* < 0.001, posthoc contrast tests: BL x CSe *F*(1,8) = 48.30, *p* < 0.001; BL x omission *F*(1,8) = 4.34, *p* = 0.071. * *p* < 0.05, ** *p* < 0.01, *** *p* < 0.001

All cell-types showed an increase in neuronal activity from baseline (BL) following the reward-predictive CS+ and reward delivery (**Figure 3D-F**). It was recently shown that two types of VTA dopamine neurons are distinguished by transient versus sustained cued-reward related activity^25^. We found that CS+ induced transient or sustained cued-reward related activity was cell-type specific. Glutamate-dopamine neurons showed sustained activity, in which activity was statistically indistinguishable between CS+ and just prior to reward delivery. In contrast, transient activity was observed in glutamate-only neurons, nonglutamate-dopamine neurons, and all dopamine neurons consisting of a statistically significant reduction in CS+ induced activity just prior to reward delivery (**Supplementary Figure 2**).

Signaling related to the CS-was also cell-type specific. To match the same approach and reward-seeking behaviors to the CS+, we examined the minority of CS-trials in which mice entered the reward port (CS-hits). Nonglutamate-dopamine neurons showed a significant suppression of activity from baseline at the time of no reward delivery (**Figure 3H-F**). No other cell-types showed significant CS-related signaling changes during CS-hits. We also examined the majority of CS-trials that were correct rejections where mice did not enter the reward port (**Supplementary Figure 3**). We observed an initial nonsignificant increase in neuronal activity followed by a cell-type specific suppression of neuronal activity that was best captured using the minimum neuronal activity during baseline, CS-, and nonreward delivery. Both nonglutamate-dopamine neurons and all dopamine neurons showed a significant reduction in neuronal activity during correctly rejected CS-trials (**Supplementary Figure 3**). Glutamate-only neurons and glutamate-dopamine neurons showed no significant modulation by correctly rejected CS-cues.

To probe cell-types for reward prediction error signaling, a subset of CS+ trials resulted in reward omission (**Figure 3L**). Reward prediction error signaling was cell-type specific. In part mediated by their transient CS+ related activity patterns, nonglutamate-dopamine neurons as well as glutamate-only neurons showed significantly suppressed activity from baseline during reward omission, reflective of prediction error (**Figure 3M-F**). However, by virtue of their sustained CS+ related activity patterns, glutamate-dopamine neurons did not signal a prediction error. Further, all dopamine neurons showed variable average responses at reward omission that were statistically indistinguishable from baseline activity (**Figure 3M-F**), likely reflecting the increased signal variability by recording glutamate-dopamine neurons and nonglutamate-dopamine neurons together.

Given that VTA dopamine neurons show heterogeneous responses to aversive stimuli^26,27^, we next assessed potential cell-type specific signaling related to aversive conditions. Mice were trained to associate a CS+ with footshock delivery and a CS-with no footshock delivery (**Figure 4**). Prior research has shown this paradigm results in more fear-related freezing behavior in response to the CS+ compared to the CS-^28^. CS+ and shock-induced neuronal signaling was cell-type specific. Both glutamate-only neurons and glutamate-dopamine neurons increased neuronal activity following the CS+ and shock compared to baseline, with glutamate-dopamine neurons showing highly elevated activity (**Figure 4B-F**). Nonglutamate-dopamine neurons did not show any elevated activity from baseline but suppressed activity between the CS+ and shock (**Figure 4B-F**). To better capture the suppressed signaling of nonglutamate-dopamine neurons, we compared the minimum neuronal activity during baseline and during shock. Nonglutamate-dopamine neurons showed a significant suppression of activity during the CS+ and shock relative to baseline using this measure (**Supplementary Figure 4**). All dopamine neurons reflected intermediate activity levels between nonglutamate-dopamine neurons and glutamate-dopamine neurons that resulted in significantly increased CS+ and shock-induced activity from baseline (**Figure 4C-F**), indicating that recordings of all dopamine neurons without considering whether they co-transmit glutamate or not results in the obscuring of aversion-related dopamine subpopulation dynamics.

**Figure 4.**
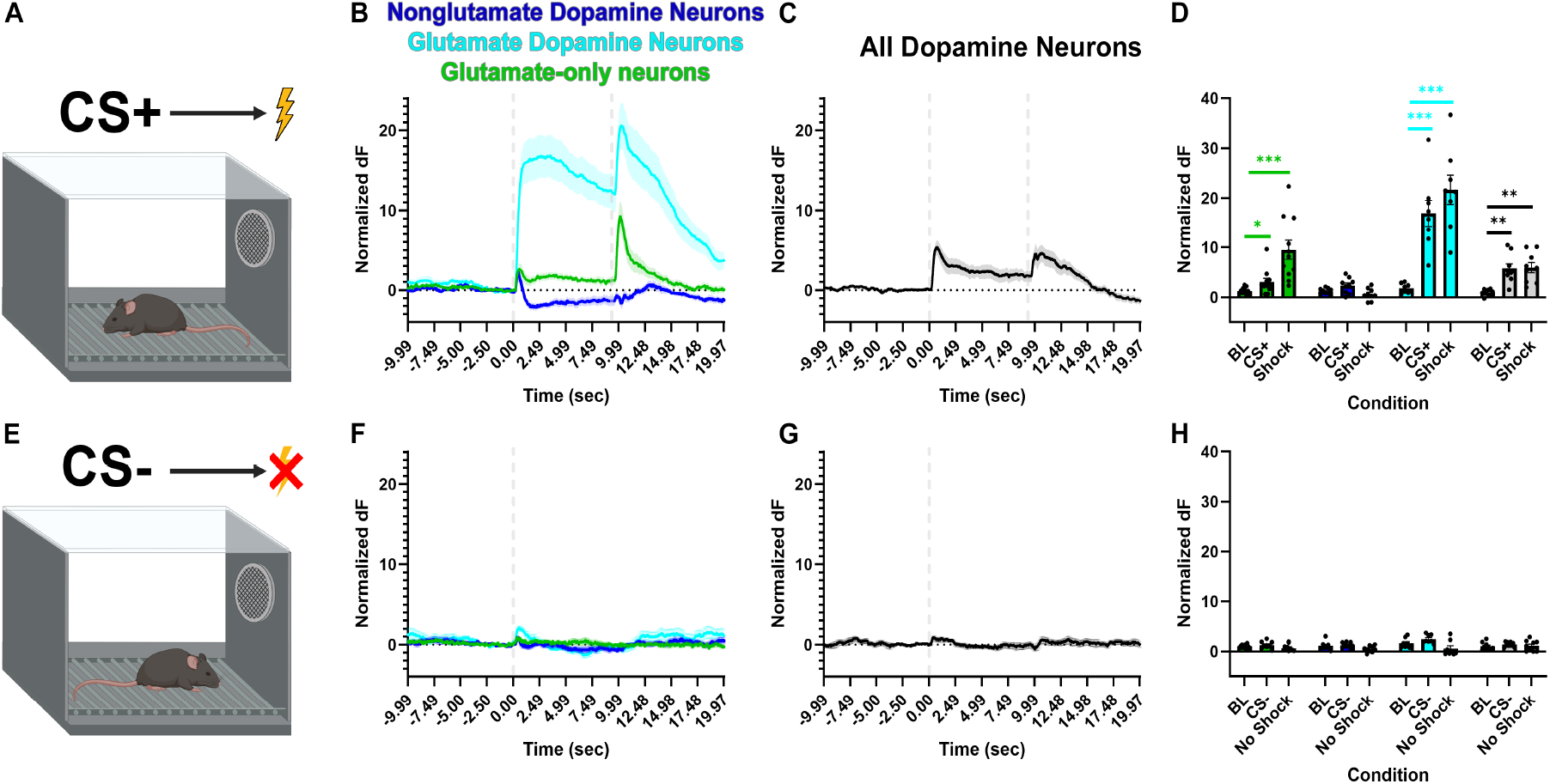
Cell-type specific cued aversive-related signaling. **A-D**. Neuronal activity during CS+ and footshock trials. Nonglutamate-dopamine (dark blue), glutamate-dopamine (cyan), glutamate-only (green), and all dopamine (black) neuronal activity. Glutamate-only neurons: main effect of epoch *F*(2,20) = 17.52, *p* < 0.001, posthoc contrast tests BL x CS+ *F*(1,10) = 6.66, *p* = 0.027; BL x shock *F*(1,10) = 22.94, *p* < 0.001. Nonglutamate-dopamine neurons: main effect of epoch *F*(2,16) = 5.96, *p* = 0.012, posthoc contrast tests BL x CS+ *F*(1,8) = 3.91, *p* = 0.083; BL x shock *F*(1,8) = 1.69, *p* = 0.230. Glutamate-dopamine neurons: main effect of epoch *F*(2,14) = 38.12, *p* < 0.001, posthoc contrast tests BL x CS+ *F*(1,7) = 33.95, *p* < 0.001; BL x shock *F*(1,7) = 48.12, *p* < 0.001. All dopamine neurons main effect of epoch: *F*(2,16) = 15.28, *p* < 0.001, posthoc contrast tests BL x CS+ *F*(1,8) = 21.48, *p* = 0.002; BL x shock *F*(1,8) = 19.23, *p* = 0.002. **E-H**. Neuronal activity during CS- and no footshock trials. Glutamate-only neurons: *F*(2,20) = 2.42, *p* = 0.114. Nonglutamate-dopamine neurons: *F*(2,16) = 2.88, *p* = 0.085. Glutamate-dopamine neurons: main effect of epoch *F*(2,14) = 5.61, *p* = 0.016, posthoc contrast tests: BL x CS-*F*(1,7) = 2.63, *p* = 0.149; BL x no shock *F*(1,7) = 2.65, *p* = 0.147. All dopamine neurons: *F*(2,16) = 0.86, *p* = 0.444. * *p* < 0.05, ** *p* < 0.01, *** *p* < 0.001

Based on our observation of cell-type specific sustained versus transient cued reward signaling, we next examined whether the footshock-predicting CS+ led to cell-type specific sustained or transient responses. Glutamate-only neuron activity was sustained between CS+ and immediately prior to footshock. However, nonglutamate-dopamine neurons, glutamate-dopamine neurons, and all dopamine neurons showed reduced changes in neuronal activity immediately prior to footshock compared with the CS+ (**Supplementary Figure 5**). No cell-type showed a significant change in activity by the CS- or time of the undelivered footshock (**Figure 4E-F**).

To assess whether VTA neurons scale their signaling of aversive events in a cell-type specific manner, mice were administered unsignaled footshocks of different intensities. Afterwards, mice were administered 0 mA footshocks accompanied by an audible click that was present during all prior footshocks (**Supplementary Figure 6**). Glutamate-only neurons were activated by all shock intensities except the 0 mA click. Nonglutamate-dopamine neurons showed no significant activation by any footshock intensity but instead showed a suppression of activity at each current intensity except the 0 mA click that was best captured using minimum activity levels (**Supplementary Figure 7**). Glutamate-dopamine neurons showed strong activation following each footshock intensity as well as the audible click during 0 mA footshocks, opposite from nonglutamate-dopamine neurons (**Supplementary Figure 6**). All dopamine neurons again reflected intermediate activity levels between nonglutamate-dopamine neurons and glutamate-dopamine neurons that resulted in significantly increased shock-induced activity from baseline across all intensities and the 0 mA audible click (**Supplementary Figure 6**).

Cell-type specific optogenetically elicited dopamine **release dynamics**

We next assessed whether activation of accumbal axons from glutamate-dopamine neurons or nonglutamate-dopamine neurons resulted in similar or different dopamine release dynamics *in vivo*. Based on VTA glutamate-dopamine neurons projecting preferentially to medial nucleus accumbens (NAcc) shell^29-31^ and VTA nonglutamate-dopamine neurons projecting preferentially to core^32^ (**Supplementary Figure 8**), VGluT2::Cre/TH::Flp mice were injected in NAcc targeting the shell/core border with an AAV encoding the dopamine sensor GRAB_DA1h_. One group of mice was also injected in VTA with an AAV encoding the red-shifted optogenetic actuator ChRmine-oScarlet in a Cre- and Flp-dependent manner to target glutamate-dopamine neurons. In a separate group of mice, ChRmine-oScarlet was delivered in a Coff/Fon manner to target nonglutamate-dopamine neurons (**Figure 5A**). ChRmine stimulation of accumbal nonglutamate-dopamine axons significantly increased the magnitude of dopamine release at every stimulation pulse compared with baseline levels (**Figure 5B-F**). The magnitude of nonglutamate axon dopamine release was significantly larger at 10 pulses compared to a single pulse. We next measured the duration of the ChRmine-evoked dopamine release by identifying the time at which the magnitude of dopamine release decreased by 50%, termed the half maximum. Nonglutamate-dopamine axons showed longer durations of dopamine release at the highest pulse numbers compared with the lowest pulse numbers, such that both 1 pulse and 5 pulses elicited shorter half maximum durations than 20 and 30 pulses (**Figure 5D**).

**Figure 5.**
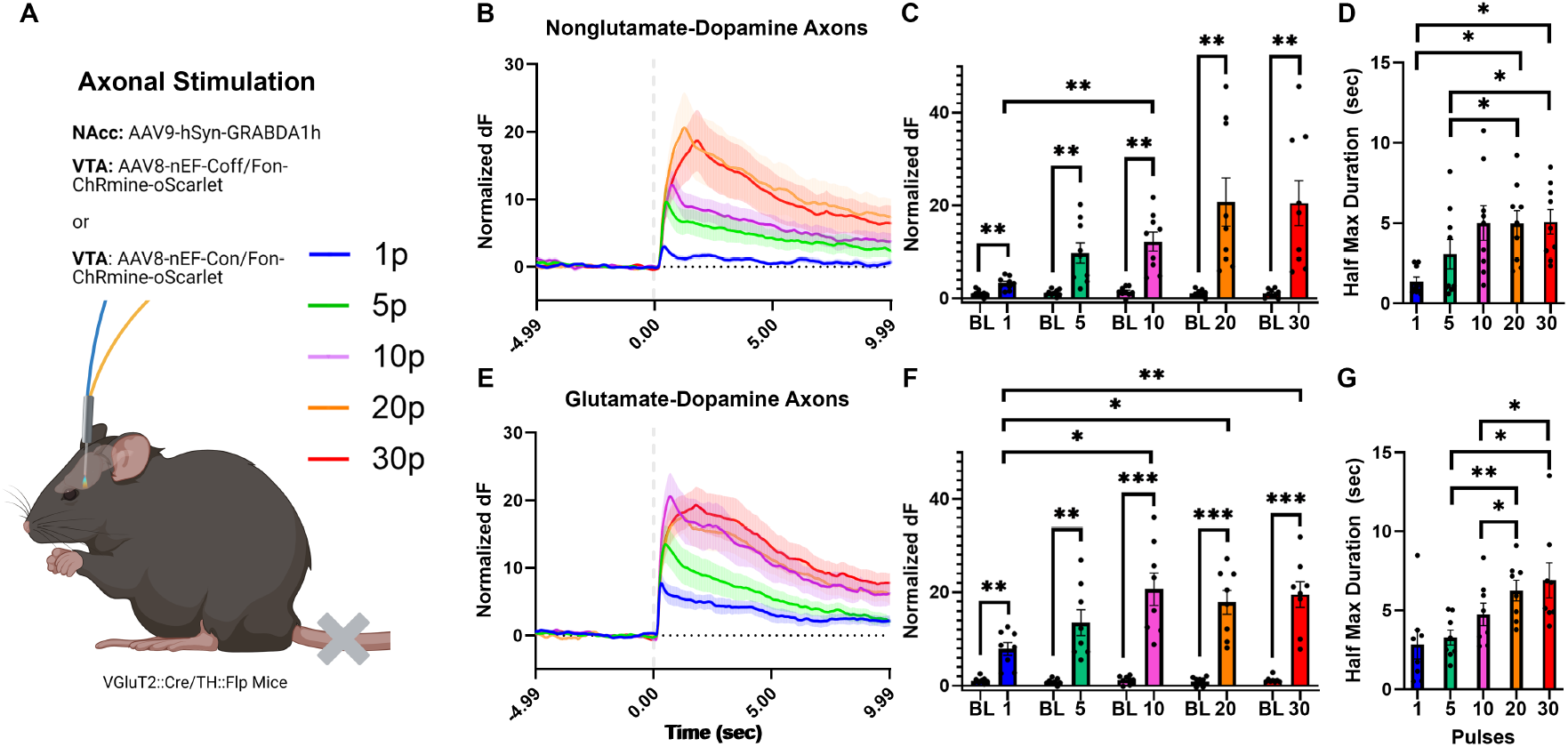
Cell-type specific *in vivo* mesolimbic dopamine release dynamics. **A**. Experimental setup: VGluT2::Cre/TH::Flp mice were injected in VTA with AAVs encoding ChRmine-oScarlet targeting nonglutamate-dopamine neurons (Coff/Fon) or glutamate-dopamine neurons (Con/Fon). The dopamine sensor GRAB_DA1h_ was expressed in NAcc and an optic fiber was implanted in the same location. Cell-type specific accumbal axons were activated with 589 nm light between 1 and 30 pulses (5 ms). **B-D**. Nonglutamate-dopamine axon ChRmine-evoked dopamine release dynamics (B), phasic magnitude of dopamine release (C), and duration of release (D, half maximum). **C**. Main effect of pulse *F*(4,32) = 10.20, *p* = 0.008, posthoc contrast tests: 1 × 10 pulses *p* = 0.008. Pulse x time interaction *F*(4,32) = 10.76, *p* = 0.006, posthoc contrast tests: 1 pulse *p* = 0.002, 5 pulses *p* = 0.005, 10 pulses *p* = 0.001, 20 pulses *p* = 0.005, 30 pulses *p* = 0.004. **D**. Main effect of pulse *F*(4,32) = 10.25, *p* < 0.001, posthoc contrast tests: 1 × 20 pulses *p* = 0.025, 1 × 30 pulses *p* = 0.012, 5 × 20 pulses *p* = 0.046, 5 × 30 pulses *p* = 0.025. **E-F**. Glutamate-dopamine axon ChRmine-evoked dopamine release dynamics (E), phasic magnitude of dopamine release (F), and duration of release (G, half maximum). **F**. Main effect of pulse *F*(4,28) = 11.47, *p* < 0.001, posthoc contrast tests: 1 × 10 pulses *p* = 0.010, 1 × 20 pulses *p* = 0.036, 1 × 30 pulses *p* = 0.006. Pulse x time interaction *F*(4,28) = 11.78, *p* < 0.001, posthoc contrast tests: 1 pulse *p* = 0.002, 5 pulses *p* = 0.003, 10 pulses *p* < 0.001, 20 pulses *p* < 0.001, 30 pulses *p* < 0.001. **G**. Main effect of pulse *F*(4,28) = 12.71, *p* < 0.001, posthoc contrast tests: 5 × 20 pulses *p* = 0.001, 5 × 30 pulses *p* = 0.020, 10 × 20 pulses *p* = 0.015, 10 × 30 pulses *p* = 0.049. NAcc = nucleus accumbens, VTA = ventral tegmental area. * *p* < 0.05, ** *p* < 0.01, *** *p* < 0.001

ChRmine stimulation of accumbal glutamate-dopamine axons significantly increased the magnitude of dopamine release at every stimulation pulse compared with baseline levels (**Figure 5E-F**). The magnitude of dopamine release following ChRmine activation of glutamate-dopamine axons was significantly higher at 10, 20, and 30 pulses compared to a single pulse. Glutamate-dopamine axons showed longer half maximum durations of dopamine release at higher pulse numbers compared with lower pulse numbers, such that both 5 and 10 pulses elicited shorter half maximum durations than 20 and 30 pulses (**Figure 5G**). Together, both nonglutamate-dopamine and glutamate-dopamine axons release dopamine into NAcc following their activation. However, glutamate-dopamine axons were more discriminative of shorter versus longer pulse trains in phasic magnitude, whereas nonglutamate-dopamine axons were more discriminative of shorter versus longer pulse trains tonically in half maximum duration.

### Neurotransmitter-specific behavioral roles of glutamate and dopamine from glutamate-dopamine co-**transmitting neurons**

Given that glutamate-dopamine neurons increased neuronal activity following both rewarding and aversive events and their learned predictors, we next aimed to determine the behavioral functions of dopamine and glutamate release from glutamate-dopamine neurons. We used Cre- or Flp-inducible AAVs encoding shRNA and GFP^33-36^ to manipulate the molecular synthesis or release machineries of dopamine (TH shRNA), glutamate (VGluT2 shRNA), or GABA (VGaT shRNA), and compared to a scrambled control sequence (scramble shRNA) in VTA neurons (**Supplementary Figure 9**). To test effectiveness and specificity, we injected the Cre-dependent shRNA vectors in VTA of VGluT2::Cre mice and after four weeks examined VGluT2 mRNA, VGaT mRNA, and TH-IR in GFP-expressing VTA neurons. In scrambled-sequence shRNA control mice, GFP neurons expressed VGluT2 (97.3 ± 0.8%), about one-fifth of which expressed VGaT (19.5 ± 1.2%) or TH (21.2 ± 1.2%). In VGluT2 shRNA mice, GFP neurons expressed VGluT2 in only 4.9% ± 0.7% of neurons while VGaT or TH expression remained about 20% each, similar to scramble controls. In VGaT shRNA mice, GFP neurons expressed VGluT2 (96.6 ± 1.5%), about one-fifth of which expressed TH (20.3 ± 0.1%), but VGaT was only detected in 2.8 ± 0.3% of neurons. In TH shRNA mice, GFP neurons expressed VGluT2 (98.6 ± 0.2%), about one-fifth of which expressed VGaT (17 ± 0.7%), but TH was nearly absent (0.1 ± 0.1% of neurons). These results demonstrate the successful knockdown of intended neurotransmission-specific transcripts in VTA VGluT2 neurons, without affecting untargeted transcripts of co-transmitters.

We next tested the role of dopamine or glutamate release machineries from VTA glutamate-dopamine neurons by injecting VGluT2::Cre mice in VTA with AAVs encoding Cre-dependent TH shRNA (to knock down TH from glutamate neurons), in TH::Flp mice AAVs encoding Flp-dependent VGluT2 shRNA (to knock down VGluT2 from dopamine neurons), or in VGluT2::Cre mice a Cre-dependent scrambled-sequence shRNA control (**Figure 6**). Mice were trained on a cued Pavlovian reward task where a CS+ predicted sucrose reward and a CS-predicted no sucrose delivery. TH knockdown from VTA glutamate neurons had no effect on reward port entries during the CS+ (**Figure 6B**) but significantly increased unrewarded entries into the reward port during the CS-within the second week of training (**Figure 6C**). VGluT2 knockdown from VTA dopamine neurons had no effect on reward port entries during the CS+ or CS-(**Figure 6E-F**). However, VGluT2 knockdown slowed the latency to enter the reward port during CS+ trials relative to scramble controls during the second week of training (**Figure 6D**). TH knockdown from VTA glutamate neurons had no effect on CS+ reward port latency (**Figure 6G**).

**Figure 6.**
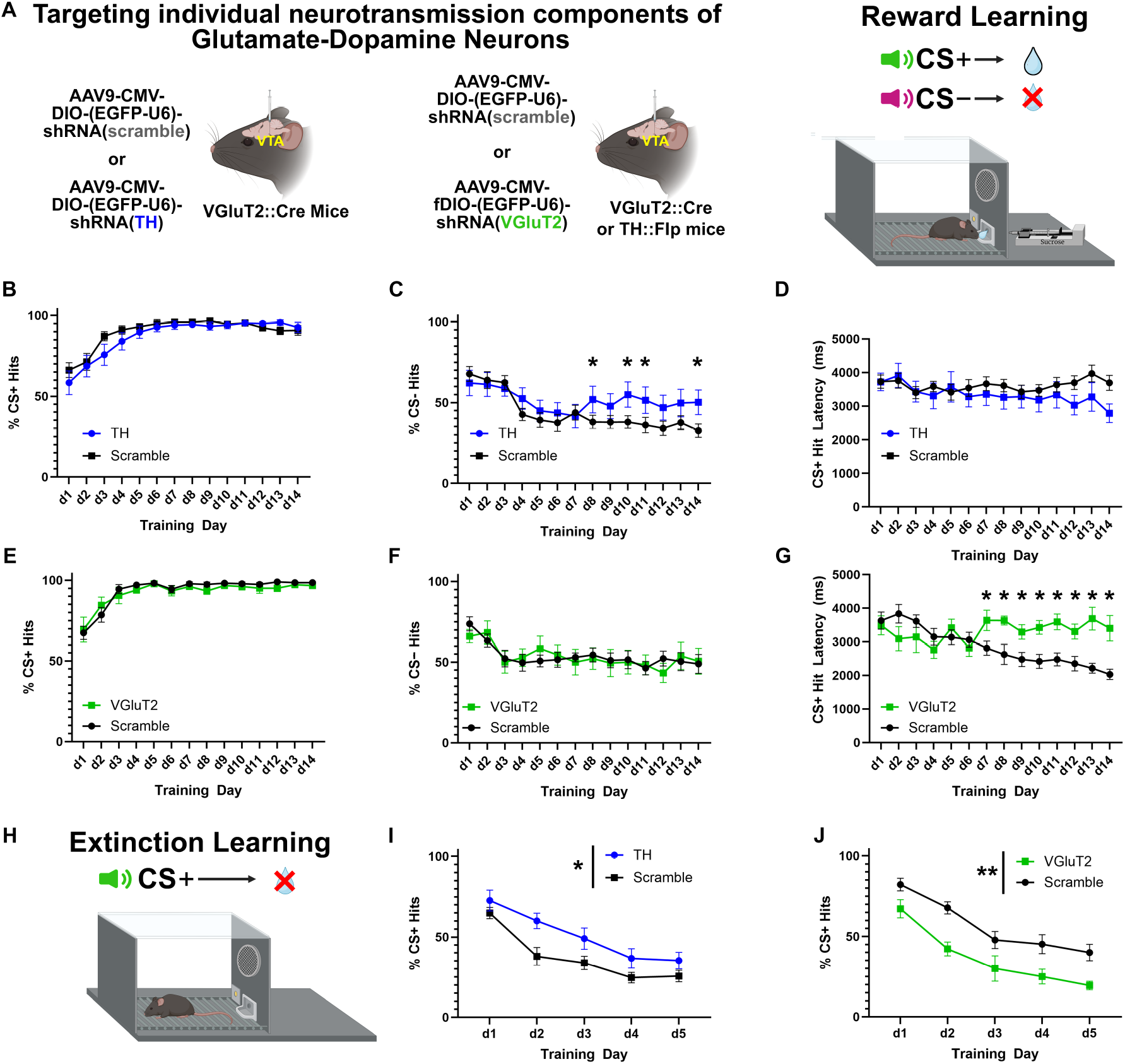
Neurotransmitter-specific knockdown from VTA glutamate-dopamine neurons alters reward-related learning and extinction of conditioned reward-seeking. **A**. Experimental setup: To test the role of dopamine in glutamate-dopamine neurons, VGluT2::Cre mice were injected in VTA with AAVs encoding Cre-dependent TH shRNA or scramble control shRNA. To test the role of glutamate in glutamate-dopamine neurons, TH::Flp mice were injected in VTA with AAVs encoding Flp-dependent VGluT2 shRNA and VGluT2::Cre mice received Cre-dependent scramble control. Mice were then trained to associate a CS+ cue with reward delivery and a CS-cue with no reward delivery. **B-D**. TH knockdown compared to scramble controls. **B**. No effect on CS+ reward port entries (day x group interaction *F*(13,260) = 0.88, *p* = 0.574). **C**. Impaired learning to reduce CS-reward port entries in second week. Day x group interaction *F*(13,260) = 3.02, *p* < 0.001, Sidak pairwise comparison day 8 *p* = 0.039, day 9 = 0.069, day 10 = 0.021, day 11 = 0.029, day 12 = 0.077, day 13 = 0.066, day 14 = 0.040. **D**. No effect on latency to enter reward port during CS+ trials (day x group interaction *F*(13,260) = 1.64, *p* = 0.139). **E-F**. VGluT2 knockdown compared to scramble controls. **E**. No effect on CS+ reward port entries (day x group interaction: *F*(13,169) = 0.62, *p* = 0.596). **F**. No effect on CS-reward port entries (day x group interaction: *F*(13,169) = 0.62, *p* = 0.711). **G**. Impaired latency to enter reward port during CS+ trials (day x group interaction: *F*(13,169) = 8.70, *p* < 0.001, all Sidak pairwise comparisons days 7-14 *p* < 0.05). **H**. Mice were trained to associate the CS+ with no reward delivery over five extinction sessions. **I**. Impaired extinction learning in TH knockdown mice (main effect of group *F*(1,21) = 5.50, *p* = 0.029). **J**. Enhanced extinction learning in VGluT2 knockdown mice (main effect of group *F*(1,13) = 13.29, *p* = 0.003). * *p* < 0.05, ** *p* < 0.01

Mice then began extinction training where the CS+ resulted in no sucrose delivery (**Figure 6H**). TH knockdown from VTA glutamate neurons slowed learning to extinguish reward port entries during the CS+ over days (**Figure 6I**). In contrast, VGluT2 knockdown from VTA dopamine neurons enhanced extinction in response to the CS+ over days compared to scramble controls (**Figure 6K**). To identify whether learning-related differences were influenced by changes in reward value, mice consumed free sucrose for one hour. Neither TH knockdown from VTA glutamate neurons nor VGluT2 knockdown from VTA dopamine neurons influenced free sucrose consumption compared with scramble controls (**Supplementary Figure 10A-F**). To determine whether differences in CS+ hits or CS+ latency were due to differences in baseline locomotor activity, mice explored an open field for 15 min. TH knockdown from VTA glutamate neurons had no effect on distance traveled while VGluT2 knockdown from VTA dopamine neurons resulted in significantly less distance traveled compared to scramble controls (**Supplementary Figure 10C-F**). We additionally measured the total time spent immobile and the number of immobile episodes in the open field assay. VGluT2 knockdown mice spent significantly more time immobile and had more immobile episodes than scramble control mice (**Supplementary Figure 10E-F**). To determine if VGluT2 knockdown led to motor abnormalities, we recorded ambulatory paw sequences and time to cross a Plexiglass catwalk. VGluT2 knockdown mice had no deficits in paw sequence regularity index nor time to cross relative to scramble controls (**Supplementary Figure 10G-F**). We also examined the performance of VGluT2 knockdown mice in the looming disc paradigm to assess locomotor performance in response to an aversive stimulus. VGluT2 knockdown mice did not show differences in latency to enter the nest, latency to freeze, or number of trials resulting in freezing, escaping, or freezing then escaping compared with scramble controls (**Supplementary Figure 10J-F**).

Mice then trained on a cued Pavlovian aversion task where a CS+ predicted footshock, a CS-predicted no footshock, and mice were examined for cue-induced fear (freezing) (**Figure 7**). TH knockdown from VTA glutamate neurons decreased CS+ induced freezing and had no effect on CS-induced freezing (**Figure 7B-F**). VGluT2 knockdown from VTA dopamine neurons had no effect on CS+ or CS-induced freezing (**Figure 7D-F**). Mice then began extinction training where the CS+ as well as the CS-resulted in no footshock (**Figure 7F**). Despite less CS+ induced freezing during acquisition, we found that extinction of CS+ induced freezing was impaired in TH knockdown mice relative to scramble controls (**Figure 7G-F**). Sidak-adjusted pairwise comparisons showed that on the first day of extinction scramble controls continued to freeze more than TH knockdown mice. However, on day 2 there was no difference in freezing between groups as control mice began to demonstrate extinction learning. During the next two extinction training days, scramble controls froze less than TH knockdown mice as they continued to extinguish CS+ induced fear, in contrast to TH knockdown mice that did not demonstrate extinction learning. There was no difference between groups for freezing over days in response to the CS-. In contrast to TH knockdown, VGluT2 knockdown from VTA dopamine neurons had no effect on CS+ or CS-induced freezing during extinction (**Figure 7I-F**). To determine whether conditioned fear-related differences in TH knockdown mice were influenced by changes in pain sensitivity, we tested mice on a hot water tail immersion assay. TH knockdown from VTA glutamate neurons did not affect pain sensitivity compared to scramble controls (**Supplementary Figure 10M-F**). To assess whether the conditioned fear and extinction-related differences in TH knockdown mice were influenced by associative learning, we assessed TH knockdown mice for nonassociative novel object learning. TH knockdown mice showed no differences in discrimination for novel objects compared to scramble controls (**Supplementary Figure 10O**). Together, glutamate and dopamine release from glutamate-dopamine neurons have dissociable contributions toward reward- and aversion-based learning and performance without affecting sensitivity to aversive events or inducing motor deficits.

**Figure 7.**
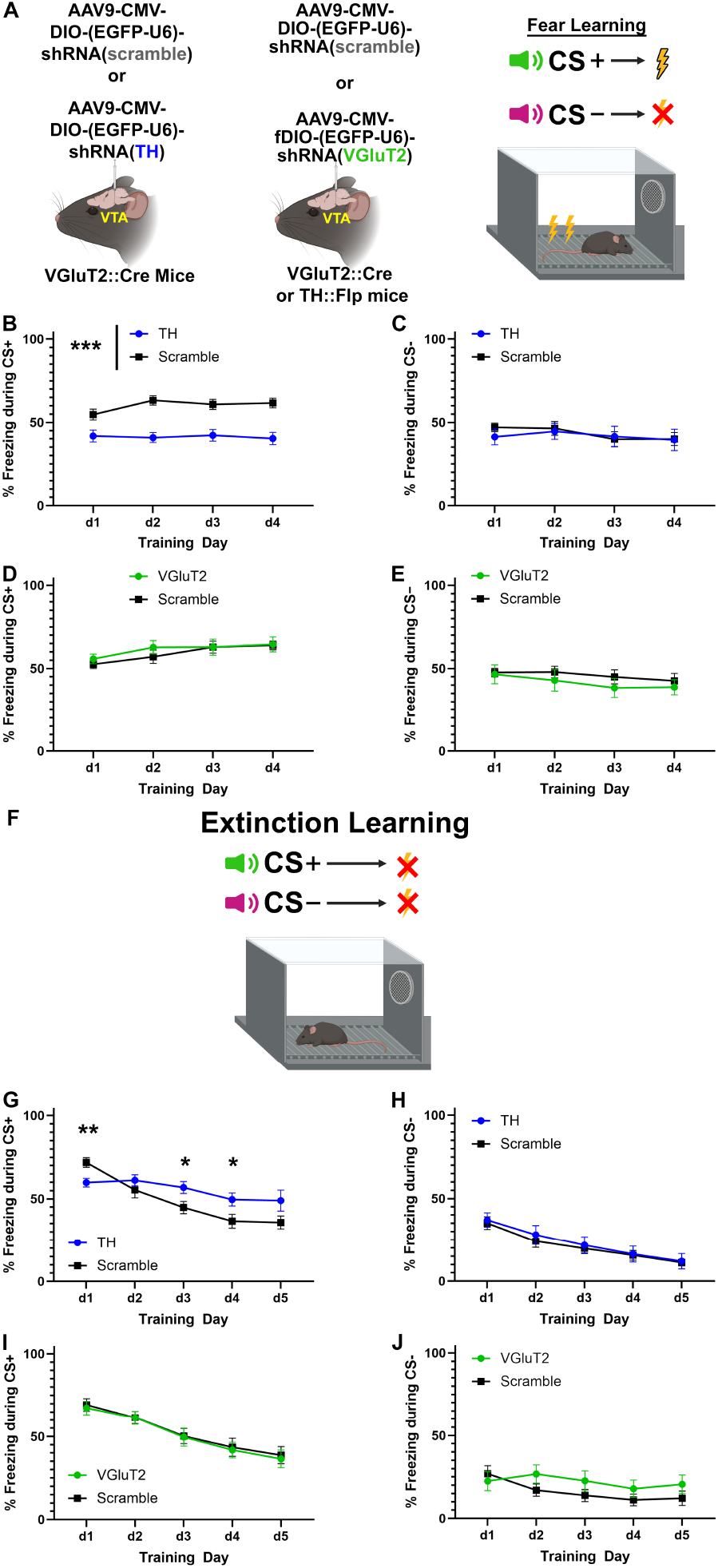
Neurotransmitter-specific knockdown from VTA glutamate-dopamine neurons alters conditioned fear learning and extinction of conditioned fear. **A**. Experimental setup: To test the role of dopamine in glutamate-dopamine neurons, VGluT2::Cre mice were injected in VTA with AAVs encoding Cre-dependent TH shRNA or scramble control shRNA. To test the role of glutamate in glutamate-dopamine neurons, TH::Flp mice were injected in VTA with AAVs encoding Flp-dependent VGluT2 shRNA and VGluT2::Cre mice received Cre-dependent scramble control. Mice were then trained to associate a CS+ cue with footshock delivery and a CS-cue with no footshock delivery. **B-C**. TH knockdown compared to scramble controls. **B**. Impaired conditioned fear learning in TH knockdown mice (main effect of group *F*(1,21) = 24.22, *p* < 0.001). **C.** No effect on CS- related freezing. **D-E.** VGluT2 knockdown compared to scramble controls **D**. No effect on CS+ related freezing or CS-related freezing (**E**). **+F**. Mice were trained to associate the CS+ with no footshock delivery over five extinction sessions and the CS-continued to be associated with no footshock delivery. **G**. Impaired extinction learning in TH knockdown mice (day x group interaction *F*(4,84) = 8.08, *p* < 0.001, Sidak adjusted pairwise comparisons day 1 *p* = 0.006, day 2 = 0.358, day 3 = 0.031, day 4 = 0.038, day 5 = 0.076). **H**. No effect on CS-related freezing. **I-J**. No effect of VGluT2 knockdown in CS+ (I) or CS-related freezing (J). * *p* < 0.05, ** *p* < 0.01, *** *p* < 0.001

### Co-transmission and neural circuit modeling of temporal-difference learning

Modeling of dopamine neuron activity has been fundamental for the development of reinforcement learning^7,37-39^. Based on our results indicating fundamental differences between glutamate-dopamine and nonglutamate-dopamine neurons, as well as glutamate or dopamine released by glutamate-dopamine neurons in motivated behavior, we developed a neural circuit model based on the neuronal activity patterns of our GCaMP-recorded dopamine neuron subtypes and behavioral analyses. The model reflects the anatomical distinction that glutamate-dopamine neurons predominantly project to medial NAcc shell^30^ while nonglutamate-dopamine neurons target NAcc core^32^ (**Supplementary Figure 8**; **Figure 8A-F**). We then tested the consequences of removing glutamate or dopamine release from glutamate-dopamine neurons in the model. Parameters were fit so that activity trajectories of the model aligned with recorded cell-type specific GCaMP data and generated behavior quantitatively consistent with reward and shock feedback-trained control conditions (**Figure 8**; variables and parameters defined in **Supplementary Table 2**). We found that knockout conditions resulted in behavior qualitatively similar to that observed in experimental data. The model was relatively flexible, and there was a broad range of parameters that produced behavior similar to *in vivo* experiments. In this model, conditioned stimuli increase activity of both nonglutamate-dopamine and glutamate-dopamine neuronal populations whose neurotransmitters in turn drive action selection in striatal targets (**Figure 8A**). Context-dependent synaptic weights evolve in response to feedback and long-term plasticity (**Figure 8B**), eventually producing neural activity profiles similar to those recorded in our GCaMP experiments (**Figure 8C**) in response to cues predicting rewarding and aversive outcomes. Learning throughout a simulated multi-day block led to an increase in the proportion of trials per day that resulted in CS+ hits or freezing in the rewarding or aversive contexts, respectively (**Figure 8D**), as well as a decrease in the latency to responding (**Figure 8E**). We found that removing glutamate from glutamate-dopamine neurons resulted in a reduction in reaction time to the reward-predicting CS+ without affecting CS+ guided reward-seeking (CS+ hits). Further, removing dopamine from glutamate-dopamine neurons resulted in reduced freezing in response to the shock-predicting CS+. Together, our model recapitulates both neural recording and behavioral trends using a simple neural circuit model which extends classic temporal-difference learning models^39^ to consider unique contributions of distinct signaling molecules released by glutamate-dopamine co-transmitting VTA populations.

**Figure 8.**
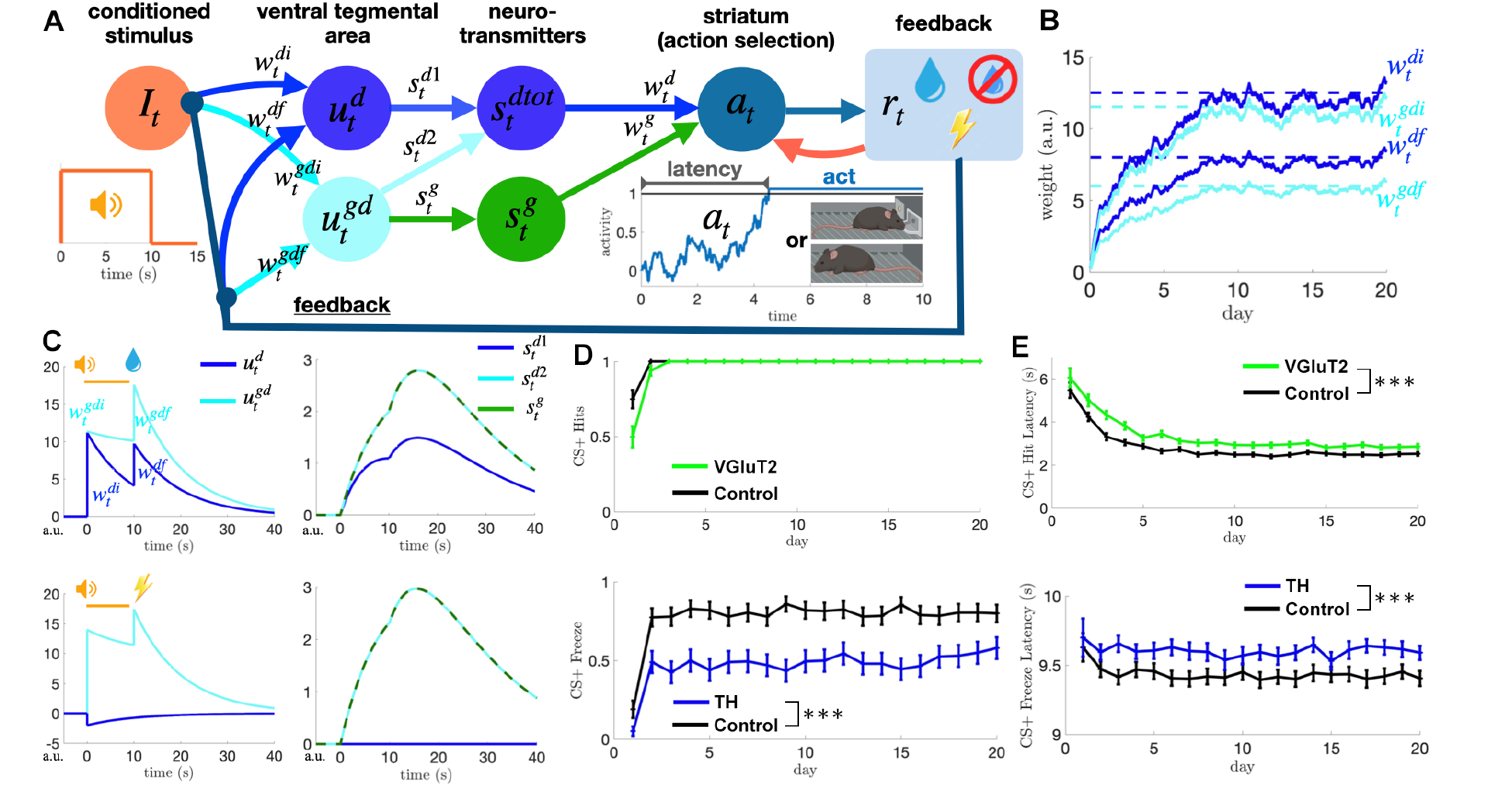
Reinforcement learning of reward and shock in a neural circuit model of action selection. **A**. Neural circuit with reward-based learning. See **Supplementary Table 2** for variable/parameter descriptions. The conditioned stimulus activates both dopamine expressing and glutamate-dopamine co-expressing neural populations in the ventral tegmental area (VTA) via weights whose strengths evolve in response to downstream feedback. Neural population activity produces neurotransmitters dopamine and glutamate which sum to drive action-selecting populations in striatum. Stronger drive leads to a more rapid rise to an action-triggering threshold, generating nose poking or freezing given prior sucrose or shock feedback. Activity must cross threshold before the conditioned stimulus terminates (10 sec) to generate an action. **B**. Synaptic weights of the input to and recurrent connections in the VTA are learned over time by probabilistic feedback (50% sucrose and 50% no sucrose) following nose pokes. **C**. Neural population and neurotransmitter activity resulting from stimulus onset (0 sec) and stimulus termination plus feedback (10 sec). Trained weights generate two peaks in both populations in the reward context but only in the co-expressing glutamate-dopamine population in the shock context. **D**. Weight training from feedback alters the fraction of trials resulting in actions each day (20 trials per day, 10 realizations), saturating to always generating nose pokes in the reward block (top) and freezing slightly more than half the time in the shock context (bottom). Removal of VGluT2 has no effect on the percent of trials in which a CS+ induced reward seeking response is made (permutation test on Euclidean distance: distance = 0.15, *p* = 0.270; 10^6^ permutations). TH removal reduces the percent of trials in which freezing occurs in response to a shock-predicting CS+ (distance = 2.54, *p* < 0.001; 10^6^ permutations). **E**. Mean latency of reward-predicting CS+ induced reward-seeking responses (top) and shock-predicting CS+ induced freezing responses (bottom) each day in each block. VGluT2 removal slows reward-predicting CS+ response latencies (distance = 1.57, *p* < 0.001; 10^6^ permutations). TH removal slows shock-predicting CS+ response latencies (distance = 0.90, *p* < 0.001; 10^6^ permutations). *** *p* < 0.001

## DISCUSSION

Despite well-established heterogeneous neuronal activity patterns^26,27,40^ and molecular diversity^14,17^ of VTA dopamine neurons, both historical and modern hypotheses on VTA dopamine function presume a homogenous influence on behavior^3,9,10,41^. A major distinction between subpopulations of VTA dopamine neurons is the presence or absence of VGluT2, the only known molecular mechanism for loading glutamate into synaptic vesicles by VTA neurons^14,15,17,19^. However, it is not understood whether molecular diversity is behaviorally relevant for motivation-related functions ascribed to VTA dopamine neurons. Further, because VTA glutamate neurons can co-transmit glutamate with dopamine or co-transmit glutamate with GABA, it has been challenging to determine the functions of VTA neurons that release *only* glutamate. By untangling glutamate-dopamine neurons from nonglutamate-dopamine neurons and glutamate-only neurons, we find that each cell-type has dissociable neuronal activity patterns and electrical properties, that glutamate-dopamine and nonglutamate-dopamine neurons have distinct dopamine release dynamics, and that dopamine and glutamate released from glutamate-dopamine co-transmitting neurons contribute to independent components of motivated behavior.

To untangle glutamate and dopamine neuron subtypes, we used INTRSECT vectors in double transgenic Cre- and Flp-expressing mouse lines. Previous research has shown the utility of INTRSECT vectors to discriminate VTA glutamate and GABA co-transmitting neurons from nonGABA-glutamate neurons and nonglutamate-GABA neurons^42^. Using VGluT2::Cre/TH::Flp mice, we segregated glutamate-dopamine neurons from nonglutamate-dopamine neurons, as previously shown^32^. Unique to the present experiments was the use of COFO vectors in a process-of-elimination strategy for selecting glutamate-only neurons as nonGABAergic nondopaminergic VTA neurons, i.e. glutamate neurons incapable of co-transmitting GABA or dopamine. Although dopamine neurons are capable of co-transmitting GABA^43^, it has not been fully investigated whether glutamate-dopamine neurons co-transmit GABA; however, stimulation of VTA glutamatergic inputs to NAcc shell that are mostly comprised of glutamate-dopamine neurons, does not result in monosynaptic activation of GABA receptors^29^. While it may be that nonglutamate-dopamine neurons are uniquely capable of GABA co-transmission, multiple mechanisms are hypothesized as critical for GABA co-transmission^43-47^ and further research will be necessary to explore each of their behavioral functions. Our initial *ex vivo* and *in vivo* characterizations identified dissociable signaling patterns of VTA glutamate-dopamine, nonglutamate-dopamine, and glutamate-only neurons. *Ex vivo*, one might predict that glutamate-dopamine neurons show intermediate electrophysiological characteristics between glutamate-only and nonglutamate-dopamine neurons. In some measures, such as firing rate and action potential duration, glutamate-dopamine neurons showed intermediate levels between the broad action potentials and slower firing rates of nonglutamate-dopamine neurons and the fast action potentials and elevated firing rates of glutamate-only neurons. However, in other measures, glutamate-dopamine neurons were more similar to either nonglutamate-dopamine neurons or to glutamate-only neurons, but not the other. For example, both glutamate-dopamine neurons and nonglutamate-dopamine neurons largely lacked rebound firing following hyperpolarization, whereas glutamate-only neurons reliably showed rebound firing. In some measures, glutamate-dopamine neurons were distinct from both glutamate-only neurons and nonglutamate-dopamine neurons. For instance, we surprisingly observed that a subset of glutamate-only neurons exhibited large I_h_ similar to nonglutamate-dopamine neurons, while glutamate-dopamine neurons largely lacked I_h_. On the whole, glutamate-dopamine neurons appear to have unique physiological properties from both glutamate-only and nonglutamate-dopamine neurons rather than a completely intermediate phenotype. Further, the identification of a subset of glutamate-only neurons with traditional electrophysiological markers of dopamine neurons such as I_h_ and D2-receptor inhibition supports prior research indicating that physiological criteria alone for VTA dopamine neuron identification are unreliable^23,48,49^.

*In vivo*, VTA glutamate-dopamine, nonglutamate-dopamine, and glutamate-only neurons showed dissociable signaling patterns related to motivationally relevant outcomes and learned cues that predicted them. During Pavlovian reward, glutamate-dopamine neurons showed sustained signaling between the CS+ and reward while nonglutamate-dopamine neurons showed transient signaling consisting of activity returning to baseline between CS+ and reward peaks. Sustained Pavlovian reward-related signaling patterns of VTA dopamine neurons were previously observed in medial VTA where glutamate-dopamine neurons predominate, and transient Pavlovian reward-related signaling patterns of VTA dopamine neurons were observed in lateral VTA where nonglutamate-dopamine neurons predominate^25^. Further, it has been found that dopamine concentration remains elevated for the entirety of a reward-predictive cue exclusively in NAcc shell where glutamate-dopamine neurons preferentially project, and not in NAcc core where nonglutamate-dopamine axons predominate^50^. The transient activity pattern of nonglutamate-dopamine neurons likely allowed for their reward prediction error signaling when an expected reward was omitted. Vice versa, the sustained activity pattern of glutamate-dopamine neurons likely prevented the possibility of producing a reward prediction error. Parallel signaling by sustained and transiently activated dopamine neurons has been hypothesized to enable proportional-differential encoding^25^. Integrating our results with this hypothesis, sustained activity in glutamate-dopamine neurons may support a behavioral state such as outcome expectation while transient activity in nonglutamate-dopamine neurons may support rate-of-change processing, together which support an ability to change expectations based on the presence or absence of the expected outcome.

Glutamate-only neurons were comparatively less activated by reward-related stimuli than dopamine neuron subtypes. However, glutamate-only neurons showed both transient activity between CS+ and reward as well as prediction error when reward was omitted, which had not been observed in prior examinations of VTA glutamate neurons^28,42^. Electrophysiologically, glutamate-only neurons exhibited more hyperpolarized resting membrane potentials, faster firing rates, and reliable rebound firing patterns following hyperpolarization compared with the dopamine subtypes. Unique subsets of glutamate-only neurons showed I_h_ and/or D2 receptor sensitivity traditionally thought to reflect a dopaminergic phenotype. We interpret these data to reflect that 1) glutamate-only neurons are a unique subset from glutamate-dopamine, glutamate-GABA, and all glutamate neurons together, and 2) glutamate-only neurons exhibit variances that may be explained by additional cell-subtypes within them. Some potential glutamate-only cell-subtypes may involve those that project locally to VTA dopamine neurons^51^ and those that express the *mu* opioid receptor^52^. During the same Pavlovian reward task, VTA dopamine and glutamate neuron subtypes were also dissociable regarding behaviors related to the CS-. Nonglutamate-dopamine neurons were the only cell-type modulated by CS-trials but did so in unique ways depending on the animal’s behavior. When mice correctly rejected entering the reward port following CS-presentation, nonglutamate-dopamine neurons suppressed their activity immediately at the CS-. However, when mice incorrectly entered the reward port following CS-presentation, nonglutamate-dopamine neurons suppressed their activity at the time that corresponded to reward delivery in CS+ trials. Together, these data indicate that nonglutamate-dopamine neurons are capable of bidirectional signaling where reward-prediction and reward result in increased activity while nonreward-prediction and nonreward result in decreased activity. Considering their CS-related activity patterns together with their prediction error signaling, decreased activity of nonglutamate-dopamine neurons during Pavlovian reward conditioning appears to coincide with the recognition that reward is not forthcoming.

The most dissociable signaling patterns of VTA glutamate-dopamine, nonglutamate-dopamine, and glutamate-only neurons were observed in response to aversive stimuli and their predictors. During Pavlovian aversive conditioning, glutamate-dopamine neurons increased activity following the CS+ and shock while nonglutamate-dopamine neurons oppositely decreased activity. The same patterns were observed in response to scaling of uncued footshock intensities. Similar to our findings in VTA glutamate-dopamine neurons, glutamate-dopamine neurons in the substantia nigra were recently found to be activated by both rewarding and aversive stimuli but not a metric of reward prediction error^53^. The VTA aversion-related signaling patterns we identified are consistent with prior research showing that VTA neurons activated by noxious stimuli are located in ventromedial VTA^26^ where glutamate-dopamine neurons predominate while VTA neurons inhibited by noxious stimuli are located in lateral VTA^26,27^ where nonglutamate-dopamine neurons predominate. It has also been hypothesized that shock-excited VTA dopamine neurons exhibit greater excitability than shock-inhibited dopamine neurons^26^; indeed, our electrophysiological results indicate greater excitability in the shock-excited glutamate-dopamine neurons compared to the shock-inhibited nonglutamate-dopamine neurons. It was previously shown that a subset of D2 receptor-expressing VTA dopamine neurons are excited by aversive stimuli via NMDA receptor signaling^54^, suggesting the subset of D2-sensitive glutamate-dopamine neurons may also express NMDA receptors, giving rise to their aversion-related excitation. Given that glutamate-dopamine neurons preferentially innervate NAcc medial shell while nonglutamate-dopamine neurons innervate NAcc core and lateral shell, our results additionally bolster the observations that aversive stimuli uniquely excite NAcc medial shell but not other accumbal subregions^55^, and that aversive cues decrease dopamine release in NAcc core and increase dopamine release in NAcc shell^56^. Together with our results in the Pavlovian reward task, this supports prior hypotheses that the medial NAcc shell differs from the core and lateral shell in its contributions to motivated behavior^50,56-64^. Congruent with other investigations of VTA glutamate neurons^28,42,65-67^, glutamate-only neurons were activated by the shock-predicting CS+ and shock itself. Only glutamate-dopamine neurons showed modulation to the audible click associated with footshock, perhaps reflective of an association with prior footshock or enhanced excitability following repeated aversive stimuli. Sustained versus transient signaling^25^was again cell-type specific. Whereas glutamate-only neurons showed sustained activity between CS+ and prior to shock, nonglutamate-dopamine neurons showed transient activity. Glutamate-dopamine neurons showed statistically reduced activity between CS+ and shock peaks, but qualitatively were highly elevated from baseline at both time points, suggestive of a largely sustained pattern.

In addition to investigating the functions of VTA dopamine and glutamate neuron subtypes, we examined the signaling patterns of all dopamine neurons without consideration for glutamate co-transmission. The signaling patterns of all dopamine neurons during Pavlovian reward were most similar to nonglutamate-dopamine neurons. However, variability of reward prediction error signaling prevented a statistical difference between baseline and error timepoints, likely driven by opposing transiently decreased nonglutamate-dopamine neuron activity and sustained activation of glutamate-dopamine neurons. Further, while the signaling patterns of all dopamine neurons during Pavlovian aversive conditioning and uncued footshock were most similar to glutamate-dopamine neurons, the elevated activity of all dopamine neurons reflected an intermediate level between glutamate-dopamine neuron activation and nonglutamate-dopamine neuron suppression. We interpret these experiments such that 1) a primary factor distinguishing the function of dopamine neuron subtypes is their response to aversive stimuli, and 2) signals from all dopamine neurons obscure subpopulation dopamine signals from glutamate-dopamine and nonglutamate-dopamine neurons.

To further assess distinctions in VTA dopaminergic subtypes, we examined whether optogenetically elicited *in vivo* dopamine release dynamics in NAcc differed between glutamate-dopamine and nonglutamate-dopamine neurons. Axonal optogenetic activation of both subpopulations elicited an increase in NAcc GRAB_DA_-related dopamine release above baseline at all pulse train levels, and scaled tonically such that longer durations of dopamine were released in response to longer pulse trains compared to shorter pulse trains. Comparatively, glutamate-dopamine axons were more discriminative of shorter versus longer pulse trains phasically while nonglutamate-dopamine axons were more discriminative of shorter versus longer pulse trains tonically. We previously observed that glutamate-dopamine neuronal activity scales with increases in reward value tonically but not phasically^66^, suggesting their phasic and tonic signaling patterns may be behaviorally meaningful. Some variability was observed in each subpopulation’s dynamics and may reflect further diversity in genetic characteristics of glutamate-dopamine and nonglutamate-dopamine neurons^14^.

Given their unique signaling patterns, dopamine release dynamics, and capability for co-transmitting glutamate with dopamine, we next sought to identify the individual contributions of glutamate and dopamine release from glutamate-dopamine neurons towards Pavlovian guided reward- and aversion-related motivated behaviors. To do so, we used recombinase-dependent interfering RNA vectors to knock down specific neurotransmission-related machineries from VTA neurons. Loss of TH resulted in deficits related to learning about less beneficial or aversion-related outcomes. For example, in Pavlovian reward conditioning, while TH knockdown mice had no deficits in learning to enter or the latency to enter the reward port during CS+ trials, they made more CS-hits than scramble controls. During extinction learning of the previously reward-predicting CS+, loss of TH reduced extinction across days compared with controls. These results are consistent with the finding that NAcc shell, which glutamate-dopamine neurons heavily innervate, is important for suppressing actions that are less likely to result in reward attainment^68^. Further, in Pavlovian aversive conditioning, TH knockdown impaired both the acquisition of a shock-predicting CS+ and the extinction of the previously shock-predicting CS+. Loss of TH had no effect on reward value assessed during free sucrose consumption, locomotor behavior in the open field test, pain detection in the tail immersion test, or nonassociative learning of the novel object task. Consistent with the present results, loss of TH from VTA VGluT2 neurons reduces optogenetic real-time place avoidance of VTA VGluT2 projections to NAcc shell^69^. Together, we interpret these results such that dopamine release from glutamate-dopamine neurons contributes to associative learning of stimuli related to aversive or less beneficial outcomes. A possible exception to this interpretation is that loss of TH impaired extinction of the freezing response to the previously shock-predictive CS+. While the lack of shock in extinction is a presumably favorable outcome, extinction learning may be contingent upon initial learning of the cue-shock association. It is therefore possible that the impairment in initial learning influenced associative learning or expression of learned fear responses in the extinction phase.

Mice with VGluT2 knockdown from glutamate-dopamine neurons led to unique differences compared to TH knockdown mice. For example, in Pavlovian reward conditioning, while VGluT2 knockdown mice had no deficits in learning to enter the reward port in response to the CS+ or reduce entering the reward port in response to the CS-, they showed a slower latency to enter the reward port during CS+ trials during the second week of training compared with scramble controls. The reduced CS+ reaction time was not due to reduced sucrose value because VGluT2 knockdown and scramble mice had similar consumption of free sucrose. The reduced CS+ reaction time was also not due to motor deficits because VGluT2 knockdown and scramble mice had no deficits in gait coordination nor the speed with which they traversed a Plexiglas catwalk. However, loss of VGluT2 resulted in reduced exploration of an open field due to a greater number of immobile episodes as well as time spent immobile. In Pavlovian aversive conditioning, VGluT2 knockdown had no effect on the acquisition of a shock-predicting CS+ and the extinction of the previously shock-predicting CS+. However, because the behavioral response in the Pavlovian aversion task is freezing and VGluT2 knockdown mice spent a significantly greater time immobile than controls in the open field assay, we tested whether a stimulus that induces active avoidance would reveal deficits in aversion-related vigor that were not captured in Pavlovian aversive conditioning. We therefore tested VGluT2 knockdown mice in a looming disc paradigm that mimicked an imminent threat from an avian predator^70^. VGluT2 knockdown had no effect on the speed with which mice responded to the looming disc stimulus or whether the response was escaping to the nest or freezing to avoid detection by the ostensive predator. Prior research has shown that loss of VGluT2 from all VTA VGluT2 neurons reduces optogenetic self-stimulation of VTA VGluT2 projections to NAcc shell^69^, suggesting a role in the attainment of rewards. Further, in dopamine neurons, loss of VGluT2 does not affect optogenetic self-stimulation of VTA dopamine neurons^71^. Together with the present results, we interpret these findings such that glutamate release from glutamate-dopamine neurons contributes to reward- and exploration-related vigor without impacting associative learning, reward value, aversion-related vigor, or gross motor deficits. Congruent with this interpretation, VTA dopamine neurons that project to NAcc shell (i.e., glutamate-dopamine neurons) may support motivational vigor in a previously learned reward task^72^. Nevertheless, the observation that glutamate receptor antagonists block place aversion resulting from optogenetic activation of VTA glutamatergic axons in NAcc shell^29^ suggests glutamate release plays a role in aversive processes not captured in the present experiments.

Recently, dopamine release from VTA has been implicated in valence-independent signaling of salient events, in which dopamine release increases in response to both rewarding and aversive stimuli^73,74^. Our somatic calcium recordings of glutamate-dopamine neurons are consistent with salience signaling, whereas calcium recordings in nonglutamate-dopamine neurons are consistent with valence signaling, in which activity increases in response to reward and decreases in response to aversive stimuli. However, our knockdown experiments in glutamate-dopamine neurons identified dissociable roles for glutamate release and dopamine release in different aspects of motivated behavior. It is therefore simultaneously possible that somatic activity of glutamate-dopamine neurons reflects salience while their distinct co-transmitters differentially influence valence signaling. Given that accumbal dopamine release can be evoked axoaxonally independent of somatic firing patterns^75-80^ and that glutamate and dopamine are released from distinct microzones within glutamate-dopamine axons^30^, further research will be necessary to identify whether glutamate and dopamine co-transmission is differentially influenced by local circuits and correlated with separate behaviors.

Finally, we developed a neural circuit model to illustrate the potential contributions of glutamate-dopamine co-transmission to reinforcement learning and action selection based on our GCaMP recordings. In the model, a conditioned stimulus influences the activity of nonglutamate-dopamine and glutamate-dopamine VTA neurons, leading to dopamine and glutamate transmission to striatal targets. Our model assumes that actions are selected in the striatum following neuronal activity crossing a context-dependent threshold. The outcome of the action feeds back to the striatum as well as to the stimulus input circuits. Over the course of conditioning, outcome feedback trains the synaptic weights of the inputs, according to the strength of the outcome’s reinforcing drive and the probability of a reinforcing outcome. Increased synaptic weight decreases the latency to cross the threshold of action selection. After conditioning in an appetitive context, modeled nonglutamate-dopamine and glutamate-dopamine VTA neurons activate in response to the conditioned stimulus as well as the rewarding outcome. In an aversive context, only the glutamate-dopamine VTA neurons respond to the conditioned stimulus and aversive outcome. In this model, simulated VGluT2 removal did not alter the probability of responding to the CS+ but significantly increased the latency to respond. Simulated TH removal reduced both the probability of freezing responses to the CS+ and increased freeze latency. Thus, our model based on GCaMP recordings of dopamine neuron subtypes recapitulates the contributions of glutamate and dopamine release toward Pavlovian reward and aversion-based motivated behaviors. However, the model output is not a perfect recapitulation of mouse behavior, likely signifying greater complexity in the circuits than captured in the model. Further complexitylikely results from further divisions of glutamate-dopamine and nonglutamate-dopamine neuronal subtypes as well as distinct sub-circuits involving different postsynaptic targets^14,29,32,81,82^.

Based on our results, we propose a new hypothesis on VTA dopamine neuron function: that dopamine neuron signaling patterns and their roles in motivated behavior depend on whether they co-transmit dopamine with glutamate or not. This hypothesis is based on four conclusions drawn from the present experiments. First, VTA glutamate-dopamine, nonglutamate-dopamine, and glutamate-only neurons are functionally dissociable in electrophysiological properties, dopamine release dynamics, and signaling dynamics related to cued rewards, predictors of nonrewards, reward prediction error, and cued and uncued aversive stimuli. Second, glutamate-dopamine and nonglutamate-dopamine neurons oppositely signal aversive events, and examining all dopamine neurons together leads to an opaque visualization of each major dopaminergic subpopulation. Further, it is likely that a comprehensive understanding of VTA dopamine function requires accounting for molecular heterogeneity of additional subtypes. Third, the functional roles of glutamate and dopamine release from glutamate-dopamine co-transmitting neurons are dissociable in reward- and exploration-related motivational vigor, cued reward learning, and cued aversion learning. Fourth, taking into account heterogeneity in dopamine neurons and multiple signaling molecules can foster the creation of reinforcement models with greater behavioral resolution.

## ACKNOWLEDGMENTS

We thank Suzanne Fulgham for her assistance with gait analysis. Research reported in this publication was supported by the National Institute On Drug Abuse and National Institute on Mental Health of the National Institutes of Health under Award Numbers R01DA047443, R01MH130576, R01MG137472 (DHR). The content is solely the responsibility of the authors and does not necessarily represent the official views of the National Institutes of Health. This research was also supported by the Institute for Cannabis Research (State of Colorado) and ABNexus (University of Colorado) (DHR). Further support was provided by R01MH122712 (AMP). The funders had no role in study design, data collection and analysis, decision to publish, or preparation of the manuscript. GraphPad Prism, Adobe Photoshop, and BioRender. com were used to generate figures and schematics.

The authors have no financial interest to be disclosed.

## METHODS

### Animals

VGluT2-IRES::Cre mice (*Slc17a6*^*tm2*(*cre)Lowl*^/J; Jax Stock #016963), TH-2A::FlpO mice (C57BL/6N-*Th*^*tm1Awar*^/ Mmmh; RRID:MMRRC_050618-MU), and VGaT-IRES::Cre mice (*Slc32a1*^*tm2*.*1*(*cre)Lowl*^/J; Jax Stock #016962) were bred at the University of Colorado Boulder. VGluT2::Cre and VGaT::Cre mice were crossed with TH::Flp mice to produce VGluT2::Cre/TH::Flp and VGaT::Cre/TH::Flp offspring, respectively. Mice were maintained in a colony room with a 12 h light/dark cycle (lights on at 22:00 h) with *ad libitum* access to food and water. All animal procedures were performed in accordance with the National Institutes of Health Guide for the Care and Use of Laboratory Animals and approved by the University of Colorado Boulder Institutional Animal Care and Use Committee.

### Stereotactic surgery

Male and female mice (8-28 weeks of age) were anesthetized with 1-3% isoflurane and secured in a stereotactic frame (Kopf). For calcium imaging experiments, INTRSECT vectors encoding GCaMP6m were used to target select neuronal populations^69,70^. In VGluT2::Cre/TH::Flp mice (4 m, 4 f), AAV8-EF1-Con/Fon-GCaMP6m (Addgene #137119, 8.5×10^12^ titer, 400 nL) was injected into VTA (AP -3.20 mm relative to bregma, ML 0.00 mm relative to midline, DV -4.34 mm from skull surface) to target glutamate-dopamine (VGuT2+/TH+) neurons. In VGluT2::Cre/TH::Flp mice (6 male, 2 female), AAV8-EF1-Coff/Fon-GCaMP6m (Addgene 137121, 7.0×10^12^ titer) was injected into VTA (AP -3.23, ML +0.30, DV -4.34 mm; 400 nL) to target nonglutamate-dopamine (VGluT2-/TH+) neurons. In TH::Flp mice (3 male, 5 female), AAV8-EF1-Coff/Fon-GCaMP6m (Addgene 137121, 7.0×10^12^ titer) was injected into VTA (AP -3.23, ML +0.30, DV -4.34 mm; 400 nL) to target all dopamine (TH+) neurons regardless of co-transmission capabilities (i.e., without respect to Cre). In VGaT::Cre/TH::Flp mice (7 male, 4 female), AAV8-nEF-(Cre-or-Flp)-Off-GCaMP6m (COFO-GCaMP6m; Neurophotonics, 5×10^12^ titer) was injected into VTA (AP -3.20, ML 0.00, DV -4.34 mm; 350 nL) to target glutamate-only (VGaT-/TH-) neurons. Total injection volume and flow rate (100 nL/min) were controlled with a microinjection pump (Micro4; World Precision Instruments, Sarasota, FL). The syringe was left in place for an additional 10 min following injection to allow for virus diffusion, after which the syringe was slowly withdrawn to prevent spread by capillary action. Mice in photometry experiments were implanted with an optic fiber cannula (400 μm core diameter, 0.66 NA; Doric Lenses, Québec, Canada) dorsal to the VTA injection site (glutamate-dopamine and glutamate-only: AP -3.20, ML +1.00 angled at 10°, DV -4.20 mm; nonglutamate-dopamine and all dopamine: AP -3.23, ML +1.30 at 10°, DV -4.25 mm). Optic fiber cannulae were angled to avoid damaging the aqueduct. All implants were secured to the skull with screws and dental cement. Mice recovered for three weeks before experimentation.

To verify cell-type specificity of the INTRSECT vectors, male and female mice were injected with GCaMP-encoding AAVs as described above to target glutamate-dopamine neurons (*N* = 3 VGluT2::Cre/TH::Flp mice), nonglutamate-dopamine neurons (*N* = 3 VGluT2::Cre/TH::Flp mice), and glutamate-only neurons (*N* = 4 VGaT::Cre/TH::Flp mice). After three weeks, mice were perfused as described in *In situ hybridization*.

For *ex vivo* electrophysiology experiments, VGluT2::Cre/TH::Flp mice (5 male, 3 female) were injected in VTA (AP -3.2, ML 0.00, DV -4.34 mm) with Con/Fon-eYFP (Addgene 55650, 5.3×10^12^ titer, 350 nL) to target glutamate-dopamine neurons, and in the same surgery were injected in VTA (AP -3.23, ML +0.30, DV -4.34 mm) with Coff/Fon-mCherry (Addgene 137134, 4.8×10^12^ titer, 350 nL) to target nonglutamate-dopamine neurons. To target glutamate-only neurons, VGaT::Cre/TH::Flp mice (5 male, 3 female) were injected in VTA (AP -3.24, ML 0.00, DV -4.32 mm) with COFO-ChRmine-oScarlet (UNC Neurotools, 3.1×10^12^ titer, 350 nL). After three weeks, mice were shipped to George Washington University for whole-cell recordings.

For *in vivo* optical stimulation and dopamine sensor photometry experiments, male and female VGluT2::Cre/ TH::Flp mice were injected in NAcc (AP +1.28, ML + 0.75, DV -4.50 mm) with pan-neuronal AAV-hSyn-GRAB_ DA1h (Addgene #113050, 5.0×10^12^ titer, 400 nL). In the same surgery, mice received either AAV8-nEF-Con/Fon-ChRmine-oScarlet (Addgene 137159, 9.0×10^12^ titer, 400 nL) in VTA (AP -3.20, ML 0.00, DV -4.30 mm) to target glutamate-dopamine neurons (5 male, 3 female) or AAV8-nEF-Coff/Fon-ChRmine-oScarlet (Addgene 137160, 9.0×10^12^ titer, 400 nL) in VTA (AP -3.22, ML +0.30, DV -4.30 mm) to target nonglutamate-dopamine neurons (5 male, 4 female). All mice were implanted with an optic fiber cannula (400 μm core diameter, 0.66 NA; Doric Lenses, Québec, Canada) dorsal to the NAcc injection site (AP +1.28, ML +0.75, DV -4.30 mm).

Implants were secured to the skull with screws and dental cement. Mice were allowed to recover for three weeks before experimentation.

To selectively knock down neurotransmission capabilities from VTA cell-types, we used a suite of Cre- or Flp-inducible AAV2/9 vectors. Each vector encodes a recombinase-inducible EGFP along with U6 promoter upstream of a short-hairpin RNA (shRNA) that has a complementary sequence to the mRNA transcript of the target gene of interest. The control vector encoded a scrambled shRNA sequence not complementary to any transcript. Vectors were either AAV9-CMV-DIO-(U6-EGFP)-shRNA(TH), AAV9-CMV-DIO-(U6-EGFP)-shRNA(VGluT2), AAV9-CMV-DIO-(U6-EGFP)-shRNA(VGaT), or AAV9-CMV-DIO-(U6-EGFP)-shRNA(Scramble). shRNA

sequences were:

VGluT2: 5’-GGCAAAGTTATCAAGGAGAAA-3’

TH: 5’-GTGCAGCCCTACCAAGATCAA-3’

VGaT: 5’-GCATCATCGTGTTCAGCTACA-3’

Scramble: 5’-CCTAAGGTTAAGTCGCCCTCG-3’

We tested the specificity and effectiveness of Cre-dependent vectors by injecting into VTA of male and female VGluT2::Cre mice. VGluT2::Cre mice were injected with either AAV9-CMV-DIO-(U6-EGFP)-shRNA(TH) (*N* = 2, BrainVTA PT-6772), AAV9-CMV-DIO-(U6-EGFP)-shRNA(VGaT) (*N* = 3, BrainVTA PT-6765), AAV9-CMV-DIO-(U6-EGFP)-shRNA(VGluT2) (*N* = 3, BrainVTA PT-5655), or AAV9-CMV-DIO-(U6-EGFP)-shRNA(Scramble) (*N* = 4, BrainVTA PT-2644) in VTA (AP -3.20, ML 0.00, DV -4.34 mm, 1.0-2.0×10^12^ titer, 400 nL). After four weeks, tissue was collected and analyzed as described in *Histology*.

In neurotransmitter knockdown behavioral experiments, we knocked down TH from glutamate-dopamine neurons by injecting AAV9-CMV-DIO-(U6-EGFP)-shRNA(TH) (BrainVTA PT-6722, 2.0×10^12^ titer, 350 nL) in VTA (AP -3.20, ML 0.00, DV -4.34) of VGluT2::Cre mice (13 male, 11 female), or knocked down VGluT2 from glutamate-dopamine neurons by injecting AAV9-CMV-fDIO-(U6-EGFP)-shRNA(VGluT2) (BrainVTA PT-8351, 2.88×10^12^ titer, 350 nL) in VTA (AP -3.20, ML 0.00, DV -4.30) of TH::Flp mice (8 male, 9 female). As a control, we injected AAV9-CMV-DIO-(U6-EGFP)-shRNA(Scramble) (BrainVTA PT-2644, 2.0×10^12^ titer) in unilateral (AP -3.20, ML 0.00, DV -4.34; 350 nL) or bilateral VTA (AP -3.20, ML ±0.30, DV -4.34; 250 nL per hemisphere) of VGluT2::Cre mice (23 male, 21 female). Control mice were compared to manipulation mice within their respective cohorts.

### Pavlovian reward conditioning

Mice were food restricted to 85% free-feeding body weight and were brought daily to behavior chambers (Med-Associates) outfitted with a custom 3D-printed reward magazine, which can be found on www.root-lab.org/code. A syringe pump (Med-Associates) was equipped to deliver 20 µl sucrose solution to the reward magazine. To acclimate mice to the reward magazine, on the first day mice were administered 30 uncued presentations of 8% sucrose solution on a variable interval 60-second schedule. In subsequent daily sessions, mice were trained to consume 8% sucrose in a Pavlovian conditioning task. The task consisted of 30 presentations (trials) of a 10 sec conditioned stimulus (CS+: 6 kHz tone) that co-terminated with sucrose reward delivery (US: 1.5 sec, onset 8.5 sec after CS+ onset). The task also included 30 trials that presented a 10 sec conditioned stimulus (CS-: white noise) that delivered no sucrose. Trials were delivered on a variable interval 90-second schedule and cues were programmed for randomized order presentation. Reward port entries were detected by infrared beam break. If a reward port entry occurred during the CS+ or CS-, it was considered a CS+ hit or CS-hit, respectively. Reward port entries outside the CS+ or CS-were considered uncued entries. For GCaMP experiments, mice trained for a minimum of 9 days, until they reached 3 consecutive days with more than 90% CS+ hits and less than 64% CS-hits. For at least one training day, mice wore dummy tethers on their optic fiber cannula to acclimate them to the recording patch cable. In the subsequent recording session, the paradigm was the same as training with the exception that there were 41 CS+ trials, 41 CS-trials, and 11% of total trials were CS error (CSe) trials in which the CS+ was presented but did not result in sucrose delivery in order to measure the neuronal response to prediction error. For knockdown experiments, mice trained for 14 days as described. After training, knockdown mice then began five days of extinction training involving 30 CS+ presentations without sucrose delivery on a variable interval 90-second schedule.

### Pavlovian fear conditioning

*Ad libitum* fed GCaMP mice were brought for four days to behavior chambers outfitted with rod flooring which was electrically connected to a shock generator (Med-Associates). The task consisted of 10 presentations (trials) of a 10 sec conditioned stimulus (CS+: 10 kHz tone) that co-terminated with footshock delivery (US: 0.5 sec, onset 9.5 sec after CS+ onset). The task also included 10 trials that presented a 10 sec conditioned stimulus (CS-: clicking) that delivered no footshock. Trials were delivered on a variable interval 90-second schedule and cues were programmed for randomized order presentation. For GCaMP mice, on the fourth day GCaMP6m signal was recorded. For knockdown experiments, mice trained for 4 days as described. After training, knockdown mice then began five days of extinction training involving 10 CS+ presentations without footshock and 10 CS-presentations on a variable interval 90-second schedule. Knockdown mice were video recorded with ANY-Maze (Stoelting). Total seconds of freezing, defined as the absence of movement except for breathing and postural adjustments was scored from videos during the CS+ and CS-cues.

### Aversive scaling

GCaMP mice were brought on a single day of testing to behavior chambers outfitted with rod flooring which was electrically connected to a shock generator (Med-Associates). During the GCaMP6m recording, mice were administered uncued footshocks of 0.25, 0.50, 0.75, and 1.00 mA intensities (0.5 sec). The presentation of shock levels was randomized on a variable interval 60-second schedule. After the delivery of three shocks per level, the shock generator was disconnected from the rod flooring and three 0.00 mA “shocks” were delivered on a variable interval 60-second schedule. For all shocks including 0.00 mA, shock delivery was accompanied by an audible click.

### GCaMP recordings

GCaMP6m was excited at 465 nm and 405 nm (isosbestic control) with amplitude-modulated signals from two light-emitting diodes reflected off dichroic mirrors and coupled into an optic fiber. Signals from GCaMP and the isosbestic control channel were returned through the same optic fiber and acquired with a femtowatt photo-receiver (Newport, Irvine, CA), digitized at 1 kHz, and recorded by a real-time signal processor (Tucker Davis Technologies, TDT). Analysis of the recorded calcium signal was performed using custom-written MATLAB scripts (available at www.root-lab.org/code). Signals (465 nm and 405 nm) were downsampled (10x) and peri-event time histograms (PETHs) were created trial-by-trial between -10 sec to +20 sec surrounding conditioned stimulus (CS+, CS-) onsets. For each trial, data were detrended by regressing the isosbestic control signal (405 nm) on the GCaMP signal (465 nm) and then generating a predicted 405 nm signal using the linear model generated during the regression. The predicted 405 nm channel was subtracted from the 465 nm signal to remove movement, photo-bleaching, and fiber bending artifacts (dF). Baseline normalized maximum Z-scores (normalized dF) were taken from -6 to -3 sec prior to CS+ or CS-onset and maximum Z-scores were taken from 0 to 2 sec following CS+ or CS-onset^28,71^. For cued outcome analyses, such as response to reward or shock, maximum Z-score were taken 0 to 2 sec following each event onset. In cases of non-events, such as nonreward in prediction error, nonreward in CS-trials, or nonshock in CS-trials, the time of the outcome during CS+ trials was imposed and the maximum Z-scores were taken from 0 to sec following each non-event. For some analyses, minimum event Z-scores were collected between 0 and 2 sec following cue or event onset and compared to the minimum baseline (−6 to -3 sec) Z-score.

### ChRmine stimulation and GRAB_DA_ recordings

*Ad libitum* fed mice were tail restricted by gauze tape and placed inside a Plexiglas container^50^. The purpose of tail restriction was to minimize locomotor behavior that may feed back onto recorded accumbal dopamine signals but still allow for full body movement. 589 nm light (10 mW) was delivered through the recording fiber at 1, 5, 10, 20, or 40 pulses (10 ms pulse width, 20 Hz) once per minute for a total of five stimuli per pulse number condition. Pulse number was randomly ordered, and 5 min of no stimulation was given between pulse number conditions. GRAB_DA_ was excited at 465 nm with an amplitude-modulated signal from a light-emitting diode reflected off dichroic mirrors and coupled into an optic fiber. Signals from GRAB_DA_ were returned through the same optic fiber and acquired with a femtowatt photo-receiver (Newport, Irvine, CA), digitized at 1 kHz, and recorded by a real-time signal processor (Tucker Davis Technologies, TDT). Analysis of the recorded calcium signal was performed using custom-written MATLAB scripts (available at www.root-lab.org/code). 465 nm signal was downsampled (10x) and peri-event time histograms (PETHs) were created trial-by-trial between -10 sec to +20 sec surrounding the onset of stimulation trains. For each trial, data were detrended by fitting a first order regression over the entire trial and subtracting the first order fit from the 465 nm channel data (dF). To assess phasic changes in dopamine transmission, maximum baseline normalized Z-scores (normalized dF) were taken from -5 to -3 sec prior to stimulation train onset and maximum stimulation Z-scores were taken from 0 to 2 sec following stimulation train onset. To assess tonic changes in dopamine transmission, the half maximum duration was calculated by the timepoint after 0 sec at which the normalized signal returned to half of the maximum observed value induced by stimulation^51^.

### Free sucrose consumption

Mice were food restricted to 85% free-feeding body weight and were brought daily to behavior chambers (Med-Associates) outfitted with a bottle containing 8% sucrose for 60 min. Licks were determined by a lickometer (Med-Associates). Reward value was defined as the free consumption of 8% sucrose on the third day of free consumption.

### Open field locomotor behavior

*Ad libitum* fed mice were placed in an open field (Any-Box, Stoelting, 40.5 × 40.5 cm, walls 35 cm high) and video recorded at 30 Hz using ANY-Maze software for 15 min. Distance traveled, number of immobile episodes, and total immobile time were calculated (ANY-Maze). In ANY-Maze, the slider for immobility detection sensitivity was set to 65% and the minimum immobile period was 2000 ms.

### Motor assessment

*Ad libitum* fed mice were placed on a Plexiglas catwalk channel (28.5 × 4 cm, side walls 8 cm high, approximately 2 ft above the ground) and video recorded from below as they traversed across (QuickTime, 30 Hz). Gait regularity was determined by analysis of paw placement sequences using the six normal step sequence patterns (NSSPs)^72,73^. Paw placements were notated via a frame-by-frame review of catwalk video recordings. Notation began with the first step taken once all paws were in frame and ended when one of the paws reached the distal end of the catwalk. One of six NSSPs^72^ were then assigned to each paw placement sequence. For a sequence to qualify as an NSSP, the following constraints were applied: 1) A minimum of four and less than eight paw placements; 2) Half-length sequences were included where there was a clear demonstration of uniformity of gait broken by only one irregular step, except at the beginning of a trial where consistency could not be determined by <4 paw placements; 3) NSSPs could begin with any paw placement of the cycle; and 4) The regularity index was not negatively influenced if mice changed from one NSSP to another without intervening paw placements, so long as each sequence contained at least four paw placements^72^. After NSSPs were assigned to each paw sequence, the regularity index (RI) was used to quantify ambulatory coordination:

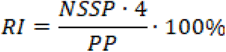

where NSSP = number of valid NSSP sequences and PP = total number of paw placements. Coordination was defined as the exclusive use of NSSPs during uninterrupted locomotion^72,73^. Thus, RI score of 100% indicates regular interlimb coordination, that no irregular steps were made, and that the trial consisted entirely of continuous sequences of NSSPs. RI score of less than 100% indicates some loss of coordination. To measure the latency to cross the catwalk, the number of frames from the start of crossing to the end of crossing was divided by the video frame rate to calculate latency time in seconds. The start of crossing was defined as the frame immediately before all paws entered the viewing plane, and the end of crossing was defined as the frame immediately before the nose reached the end of the catwalk. Mice were given 2-3 trials and only the trial with the shortest latency was scored.

### Tail immersion test

*Ad libitum* fed mice were briefly restrained in a restraint tube (Kent Scientific). Approximately 60% of distal tail was placed in 50°C heated water and the latency to remove the tail was determined. If the mouse did not remove the tail after 10 sec, the mouse was not analyzed (1 mouse).

### Novel object recognition test

*Ad libitum* fed mice were placed in an open field (AnyBox, Stoelting, 40.5 × 40.5 cm, walls 35 cm high) together with two identical copies of an object, termed the familiar object. Familiar objects were either a plastic bottle (Uxcell, 150 mL, 9 cm height x 5 cm diameter, filled with brown paper towels to decrease translucency and adorned with colored tape to increase visual complexity) or a block of interlocking plastic bricks (MEGA BLOKS, Mattel, 9.5 × 3 × 6.5 cm). Bottle and brick objects were selected for similarity in size, material, and complexity. Objects were positioned in the back left and front right corners of the apparatus 7 cm from the left or right wall and 7 cm from the back or front wall and were secured to the floor with adhesive hook and loop tape (3M). Mice were allowed to explore the familiar objects for 10 min, then were returned to the home cage for 40 min. Meanwhile, both familiar objects were removed. A third identical copy of the familiar object was placed in either the back left or front right corner. A novel object—either the bottle if the familiar object was the brick or the brick if the familiar object was the bottle—was placed in the opposite corner. The familiar object and location of the novel object were randomized across subjects. After the 40 min timeout, mice were returned to the apparatus and were allowed to explore the familiar object and the novel object for 10 min. The familiarization phase and the novel phase were tracked using ANY-Maze software, which calculated the time mice spent investigating each object. When the mouse’s head was within 25 mm and oriented toward an object, investigation was scored. Investigation was not scored when the mouse’s center point overlapped with the object in the visual field, which indicated that the animal had climbed on top of the object. The orientation angle was set to 90°. Mice were removed from analysis if the total investigation time was less than 20 sec in either stage. To determine whether mice could discriminate between the familiar and novel object, the discrimination score was calculated as:

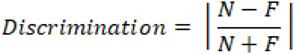

where N = time spent investigating the novel object in the novel phase, F = time spent investigating the familiar object in the novel phase, and vertical bars represent absolute value^74^.

### Looming disc

*Ad libitum* fed mice were placed in an open field (AnyBox, Stoelting, 40.5 × 40.5 cm, walls 35 cm high) together with a custom 3D printed shelter (available at www.root-lab.org/code) in the corner of the box. After at least 8 min of acclimation, when mice entered the center of trigger zone near the shelter, a looming disc was presented on an overhead monitor. The looming disc was a black disc that expanded at a diameter of 2° of visual angle to 20° in 250 ms and remained at that size for 250 ms^58^. The expanding black disc was repeated 15 times with 500 ms pauses. The trial ended 30 sec after the end of the looming disc animation. One trial per mouse was performed. From ANY-Maze-recorded videos, we analyzed the latency until mice entered the shelter and the latency until mice performed a freezing response. If no shelter entry response was made, the escape time was not examined further. If no freezing response was made, the freezing time was not examined further.

### Histology

Mice were anesthetized with isoflurane and perfused transcardially with phosphate buffer followed by 4% (w/v) paraformaldehyde in 0.1 M phosphate buffer, pH 7.3. Brains were extracted, post-fixed overnight in the same fixative and cryoprotected in 18% sucrose in phosphate buffer at 4°C. Coronal sections containing the VTA or NAcc (30 μm) were taken on a cryostat, mounted to gelatin-coated slides, and imaged for GFP or oScarlet and optical fiber cannula placement on a Zeiss Axioscope depending on experimental condition.

### In situ hybridization

Mice were anesthetized with isoflurane and perfused transcardially with phosphate buffer followed by 4% (w/v) paraformaldehyde in 0.1 M phosphate buffer, pH 7.3. VTA sections (14 μm) were mounted onto Fisher SuperFrost Plus slides and RNAscope *in situ* hybridization was performed according to the manufacturer’s instructions. Briefly, sections were treated with heat and protease digestion followed by hybridization with target probes for mouse TH (ACDBio #317621), VGluT2 (ACDBio #319171), and VGaT (ACDBio #319191). Sections were then immunolabeled with mouse anti-GFP (Clontech JL-8) and anti-mouse Alexa 488 (Jackson ImmunoSecondaries 715-545-150). Slides were coverslipped in Prolong Diamond with DAPI (ThermoFisher) and imaged on a Nikon A1R confocal (20X) with z-stacked images in 3 µm steps at 512 × 512 pixel resolution.

### Slice preparation for electrophysiology

After one week of acclimation to George Washington University, acute slices were prepared as previously described^49^ from mice anesthetized with ketamine (100 mg/kg) and dexmedetomidine (0.25 mg/kg). Mice were perfused with 34°C NMDG ringer: 92 mM NMDG, 2.5 mM KCl, 1.2 mM NaH2PO4, 30 mM NaHCO3, 20 mM HEPES, 25 mM glucose, 5 mM sodium ascorbate, 2 mM thiourea, 3 mM sodium pyruvate, 10 mM MgSO4, and 0.5 mM CaCl2^75^. Following perfusion, the brain was rapidly dissected and horizontal slices (220 mM) were prepared in warmed NMDG ringer using a vibratome. Slices recovered for 1 h at 34°C in oxygenated HEPES holding solution: 86 mM NaCl, 2.5 mM KCl, 1.2 mM NaH2PO4, 35 mM NaHCO3, 20 mM HEPES, 25 mM glucose, 5 mM sodium ascorbate, 2 mM thiourea, 3 mM sodium pyruvate, 1 mM MgSO4, and 2 mM CaCl2^75^ and then were held in the same solution at room temperature until use.

### Electrophysiology

Electrophysiological recordings were performed using a Sutterpatch integrated patch amplifier. Midbrain slices were continuously perfused at 1.5–2 ml/min with ACSF (28–32°C) containing the following: 126 mM NaCl, 21.4 mM NaHCO3, 2.5mM KCl, 1.2 mM NaH2PO4, 2.4mM CaCl2, 1.0 mM MgSO4, and 11.1 mM glucose. Potassium gluconate internal solution was used for all recordings and contained the following: 35 K-gluconate, 10 HEPES, 4 KCl, 4 Mg-ATP, 0.3 Na3-GTP. Junction potential was not corrected for. Cell-types were identified by eYFP, tdTomato, or oScarlet fluorescence. Capacitance was measured using Sutterpatch software upon breakin. Input resistance was calculated using Ohm’s law and the voltage responses to a series of hyperpolarizing current steps. Steps that induced a sag were excluded from calculation of input resistance. I_h_ was measured in voltage clamp. Cells were held at -50 mV and were subjected to a series of 500 ms hyperpolarizing voltage steps ranging from -10 to -50 mV. The I_h_ was determined by the mean current at the end of the 500 ms step subtracted from the mean leak current at the onset of the step. Cells were considered I_h_+ if they exhibited a current >25 pA in response to a 50 mV step. Spontaneous action potentials were collected in whole-cell current clamp mode. Action potential properties were measured using Sutterpatch’s action potential analysis module. Properties of between 10-50 action potentials (depending on firing rate of cell) collected shortly after break in were averaged from each cell. While bursting in slices was rare, action potentials in bursts were not included in analysis of action potential properties. Evoked action potentials were measured in response to escalating depolarizing 500 ms current injections (increments of 25 pA). Cell attached recordings were obtained in the loose-patch configuration^76^. Baseline firing rate was determined over a thirty-second period at the end of three minutes of stable firing. For recordings examining responses to D2 receptor stimulation, 1 µM quinpirole^24^ was added to the bath after a five-minute baseline of steady firing, and firing rate was again measured over a 30-second period 10 min after quinpirole addition.

### Statistical tests

GCaMP behavior and neuronal experiments, GRAB_DA_ experiments, and knockdown experiments were statistically analyzed in SPSS (IBM) or Prism (GraphPad) and figures generated in Prism and Photoshop (Adobe). For all tests, α was set to 0.05.

To compare training-related differences before GCaMP recording, mixed ANOVAs compared days of training and cell-type group for percent CS+ hits and CS-hits. Analyses of neuronal activity first consisted of repeated measures. To identify cell-type specific GCaMP signaling changes, within-subjects ANOVAs compared differences between baseline, cue, and outcome for each cell-type. Following up significant main effects of epoch, we prioritized determining differences in neuronal activity between baseline and cue, or baseline and outcome. Therefore, simple contrast tests were used. To assess transient versus sustained signaling, paired t-tests compared CS+ maximum Z-score to the pre-reward or pre-shock outcome (−2-0 sec before outcome). To assess aversion scaling, within subjects ANOVAs compared maximum Z-score 0-2 sec after shock across independent variables of current and epoch. Sidak-adjusted pairwise comparisons followed up significant main effects or interaction. For GRAB_DA_ experiments, ANOVAs compared maximum Z-score and time to half maximum Z-score following optogenetic pulses across epoch and pulse number for each cell-type specific projection. Sidak-adjusted pairwise comparisons followed up main effects or interaction. For knockdown reward or fear behavioral experiments, mixed ANOVAs compared variables of interest across days and groups. Sidak adjusted pairwise comparisons followed up significant main effects or interaction. For other knockdown comparisons between two groups alone (e.g., free sucrose licks) independent two-tailed t-tests were conducted. Where the assumption of normal distribution was not met as determined by Shapiro-Wilk test, a non-parametric two-tailed Mann-Whitney U test was used. Homogeneity of variance was confirmed in all comparisons with *F*-tests. To compare the number of trials resulting in different responses to the looming disc stimulus between knockdown and control mice, Fisher’s exact test was used. For all ANOVAs, if the assumption of sphericity was not met (Mauchley’s test), the Greenhouse-Geisser correction was used.

Electrophysiological data were analyzed using Sutterpatch software and GraphPad Prism. Repeated measures were analyzed using a two-way repeated measure ANOVA or mixed-effect model followed by Dunnett’s multiple comparison test. AP half-width, AHP size and firing rate were measured by a one-way ANOVA or Welch ANOVA (when unequal SDs were present) followed by Tukey’s test.

### Creating reinforcement learning model of VTA-striatum in reward/shock feedback task

To model neural circuit learning due to reward and shock feedback, we extended the classic temporal-difference learning framework for classical conditioning^39^ to incorporate co-expressing neural populations with slow synapses triggering an action selecting population. We now describe the evolution equations for our neural circuit model. As identified in our calcium-imaging experiments, we incorporate a population of pure dopamine expressing neurons 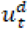 from the ventral tegmental area (VTA) and a co-expressing population 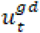 that releases glutamate and dopamine. These are each driven by the stimulus and feedback according to the differential equation (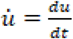: time derivative)

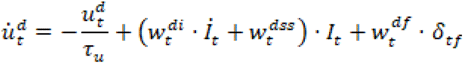

and

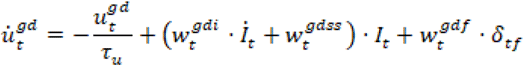

where sec is the slow decay timescale of activity in both neural populations established either by neuromodulation or slow short-term synaptic plasiticity^77^. The conditioned stimulus input *I*_*t*_=*H*_*t*_ − *H*_*tf*_ is a square wave lasting until the potential feedback time *tf* = 10 sec, generating both instantaneous 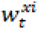 and steady state 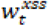 responses. Feedback *rt* = (δ_*tf*_ χ_*t*_ potentially generated at the t = *tf* is either reward χ_*t*_= χ_*rew*_

, shock χ_*t*_ = χ_*shock*_, or nothing χ_*t*_ = 0, contributing to the learning of input and feedback weights 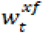 via a feedback-driven long-term plasticity rule

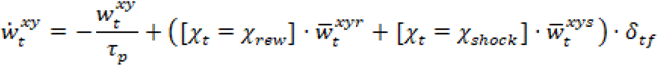

where reward versus shock feedback drives the long term weight towards different steady state values, parameterized in the reward context as 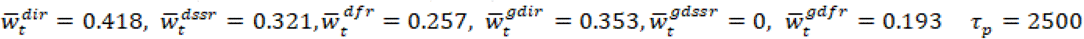, sec, and in the shock context as 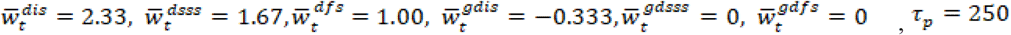, sec to qualitatively match time series of our neural population recordings. Synaptic release by these neural populations is then given by the evolution equations

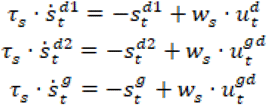

with timescales *τ*_*s*_ = 10 sec and neural population weights *w*_*s*_ = 0.3. Note, thus the total dopamine driving the action selection circuit in striatum is 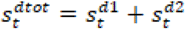. Dopamine and glutamate are then weighted and noise (*σ* = 0.224) is added to account for stochasticity so the action variable evolves as

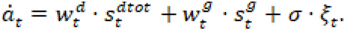

Given the learned context *c* ∈ {*χ*_*rew*,_ *χ*_*shock*_}, if ^*a*^*t* crosses the decision threshold *θ* before *t* =*tf*, then either a nose poke or freeze response is generated at the first time ^*a*^*t*≥ *θ*. A rapid learning rule akin to the long-term plasticity rule uses feedback to train the neurotransmitter weights and choice threshold according to the context so if *c* = *χrew*, then 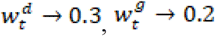, and *θ* →0.8, while for *c* = *χ*_*shock*_, then then, 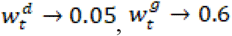, and *θ* → 8.5. For glutamate (dopamine) knockout experiments in the reward (shock) context, we set 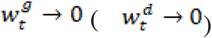.

### Testing model on numerical knockout experiments and significance measures

To simulate 10 different subjects each, we ran 10 realizations each of the four different task conditions: (1) 50/50 rewarded control; (2) rewarded glutamate knockout; (3) shock control; and (4) shock dopamine knockout. All trainable weights were initialized to zero at the beginning of a block. Each simulated day contained 20 sequential trials and 20 simulated days were run per block. Each trial lasted 40 sec with a 10 sec conditioned stimulus time and 30 sec post-choice time. Rewarded trials were drawn according to a Bernoulli process with success parameter 0.5. Evolution equations we solved using the Euler-Maruyama method with. We recorded whether a choice was made on each trial and if so the reaction time, using these to compute the mean and standard error each day across subjects for each block. The 10 subject means were then used to compute the Euclidean pairwise distance between choice fractions and reaction times between control and knockout conditions. This was compared to a distribution of distances computed from shuffled versions of the data set to compute *p*-values for these statistics in the two blocks.

### Model code availability

MATLAB code for reproducing model time series and statistical tests is available at the Github repository: https://github.com/zpkilpat/coexpressing.

## SUPPLEMENTARY INFORMATION

**Supplementary Figure 1.**
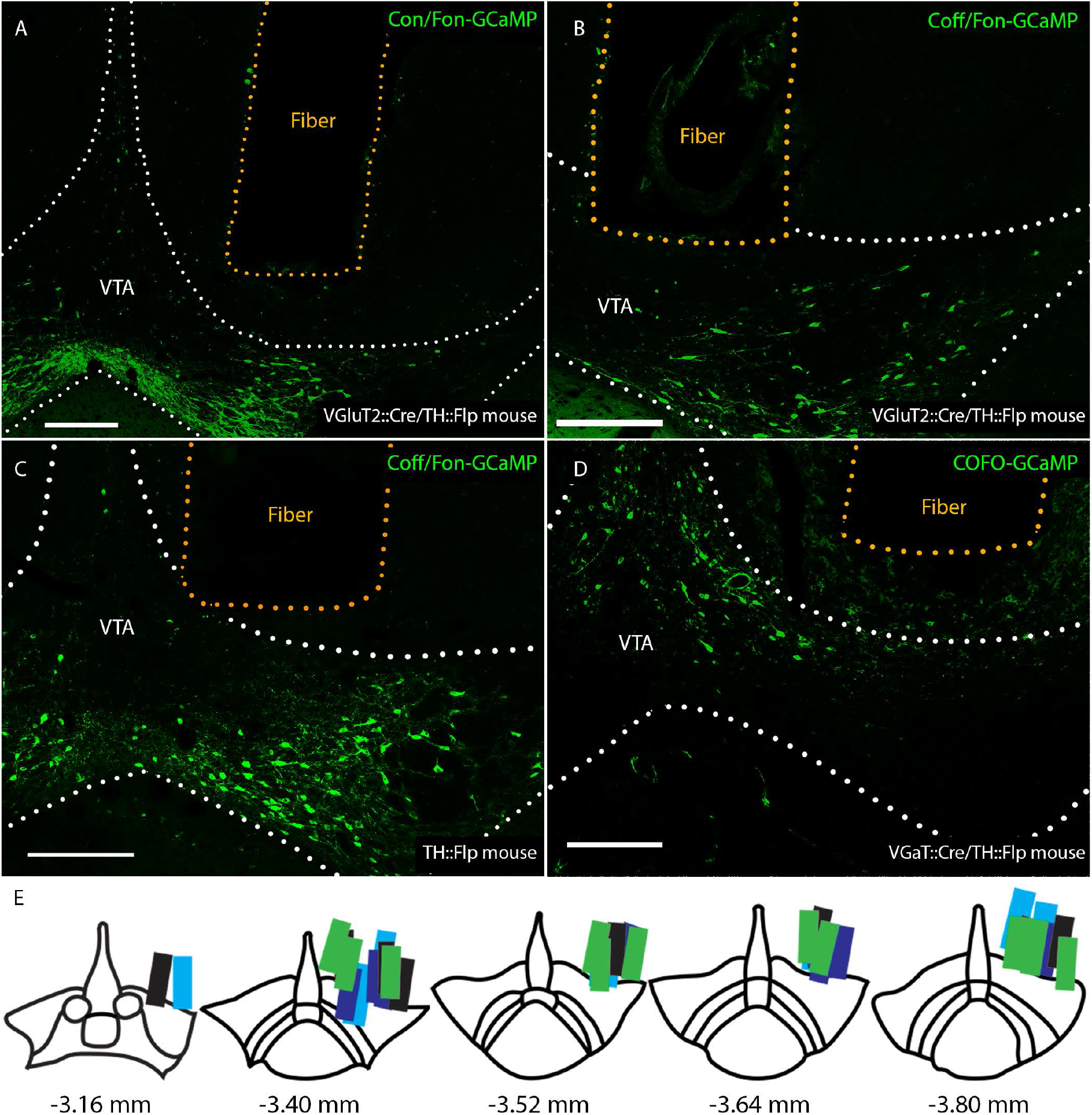
Histological validation for GCaMP experiments. Example optic recording fiber placement and viral GCaMP6m expression in VTA with (**A**) Con/Fon-GCaMP in VGluT2::Cre/TH::Flp mouse targeting glutamate-dopamine neurons, (**B**) Coff/Fon-GCaMP in VGluT2::Cre/TH::Flp mouse targeting nonglutamate-dopamine neurons, (**C**) Flp-dependent GCaMP in TH::Flp mouse targeting all dopamine neurons, and (**D**) COFO-GCaMP in VGaT::Cre/TH::Flp mouse targeting glutamate-only neurons. All scale bars = 200 µm. VTA = ventral tegmental area. (**E**) Illustration of VTA optic fiber placements for all mice in GCaMP recording experiments. Green = glutamate-only; cyan = glutamate-dopamine; dark blue = nonglutamate-dopamine; black = all dopamine. Numbers refer to anteroposterior plane from bregma.

**Supplementary Figure 2.**
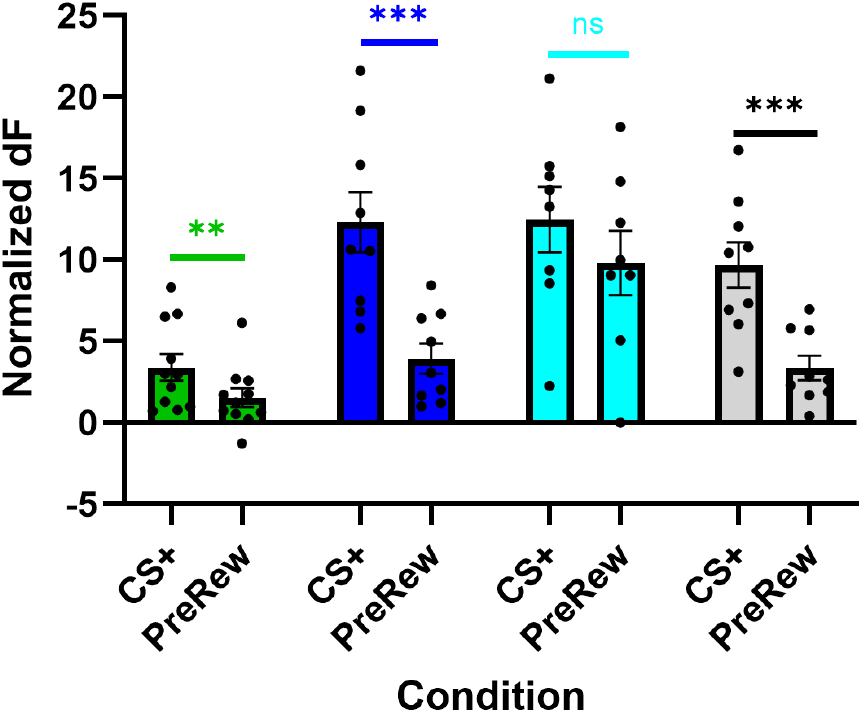
Cell-type specific transient and sustained cued reward-related signaling. Neuronal signaling in response to the CS+ and immediately before reward delivery in nonglutamate-dopamine (dark blue), glutamate-dopamine (cyan), glutamate-only (green), and all dopamine (black). Glutamate-only neurons *t*(10) = 3.71, *p* = 0.004. Nonglutamate-dopamine neurons *t*(8) = 8.47, *p* < 0.001. Glutamate-dopamine neurons *t*(7) = 1.39, *p =* 0.207. All dopamine neurons *t*(8) = 6.56, *p* < 0.001. ns = not significant, ** *p* < 0.01, *** *p* < 0.001

**Supplementary Figure 3.**
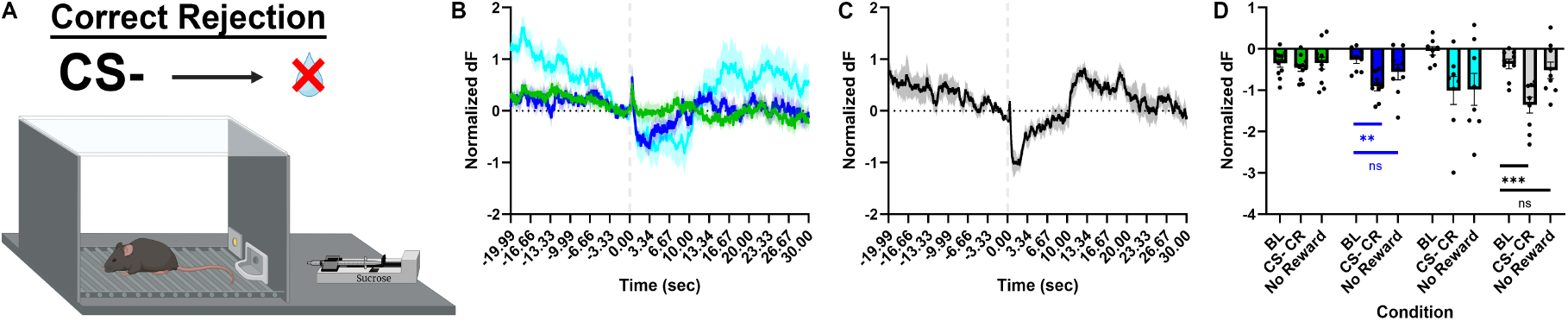
Cell-type specific correct rejection signaling. **A**. Neuronal signaling during correct rejection trials where mice did not enter the reward port during CS-trials. **B-C**. Nonglutamate-dopamine (dark blue), glutamate-dopamine (cyan), glutamate-only (green), and all dopamine (black) neuronal activity. **D**. Glutamate-only neurons: *F*(1,20) = 0.37, *p* = 0.697. Nonglutamate-dopamine neurons: main effect of epoch *F*(2,16) = 6.46, *p* = 0.009, posthoc contrast tests BL x CS-*F*(1,8) = 22.74, *p* = 0.001, BL x No Reward *F*(1,8) = 2.08, *p* = 0.187. Glutamate-dopamine neurons: *F*(2,14) = 3.47, *p* = 0.060. All dopamine neurons: main effect of epoch *F*(2,16) = 17.74, *p* < 0.001; posthoc contrast tests BL x CS-*F*(1,8) = 44.17, *p* < 0.001, BL x No Reward *F*(1,8) = 0.77, *p* = 0.405. ns = not significant, ** *p* < 0.01, *** *p* < 0.001

**Supplementary Figure 4.**
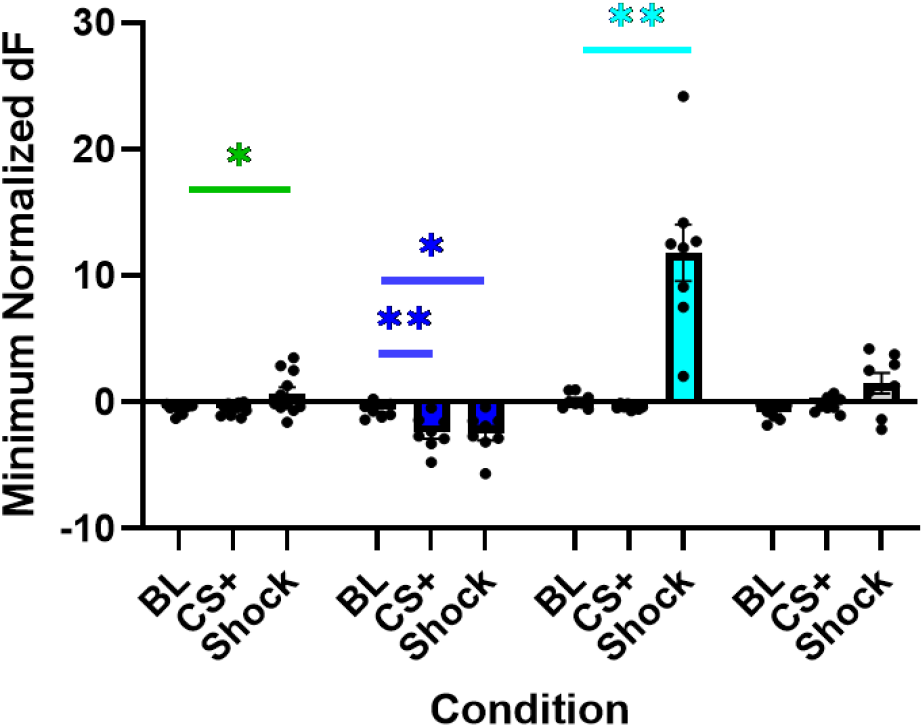
Cell-type specific suppression of neuronal activity in aversion-related signaling. Minimum normalized GCaMP response at baseline, CS+, and shock delivery in nonglutamate-dopamine (dark blue), glutamate-dopamine (cyan), glutamate-only (green), and all dopamine (black). Glutamate-only neurons: main effect of epoch *F*(2,20) = 6.73, *p* = 0.022, posthoc contrast tests BL x CS+ *F*(1,10) = 0.02, *p* = 0.878; BL x shock *F*(1,10) = 6.531, *p* = 0.029. Nonglutamate-dopamine neurons: main effect of epoch *F*(2,16) = 9.38, *p* = 0.002, posthoc contrast tests BL x CS+ *F*(1,8) = 13.71, *p* = 0.006; BL x shock *F*(1,8) = 8.36, *p* = 0.020. Glutamate-dopamine neurons: main effect of epoch *F*(2,14) = 28.26, *p* = 0.001, posthoc contrast tests BL x CS+ *F*(1,7) = 4.17, *p* = 0.080; BL x shock *F*(1,7) = 27.49, *p* = 0.001. All dopamine neurons main effect of epoch: *F*(2,16) = 4.59, *p* = 0.057, posthoc contrast tests BL x CS+ *F*(1,8) = 2.51, *p* = 0.152; BL x shock *F*(1,8) = 4.86, *p* = 0.059. * *p* < 0.05, ** *p* < 0.01, *** *p* < 0.001

**Supplementary Figure 5.**
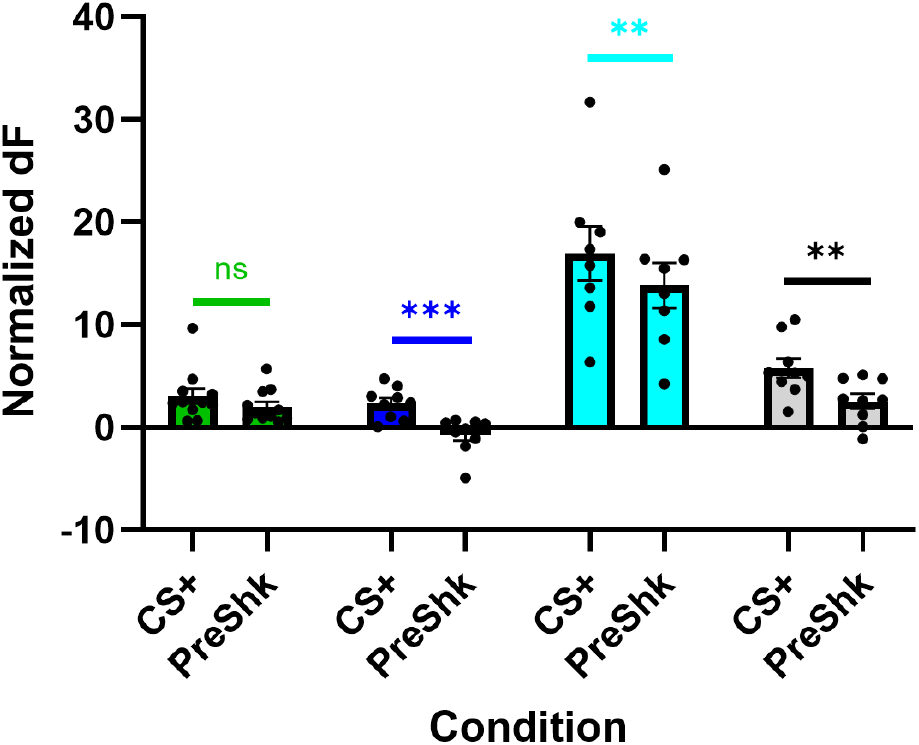
Cell-type specific transient and sustained cued aversion-related signaling. Neuronal signaling in response to the CS+ and immediately before shock delivery in nonglutamate-dopamine (dark blue), glutamate-dopamine (cyan), glutamate-only (green), and all dopamine (black). Glutamate-only neurons *t*(10) = 1.50, *p* = 0.163. Nonglutamate-dopamine neurons: *t*(8) = 5.33, *p* < 0.001. Glutamate-dopamine neurons *t*(7) = 3.64, *p* = 0.008; All dopamine neurons *t*(8) = 4.76, *p* = 0.001. ns = not significant, ** *p* < 0.01

**Supplementary Figure 6.**
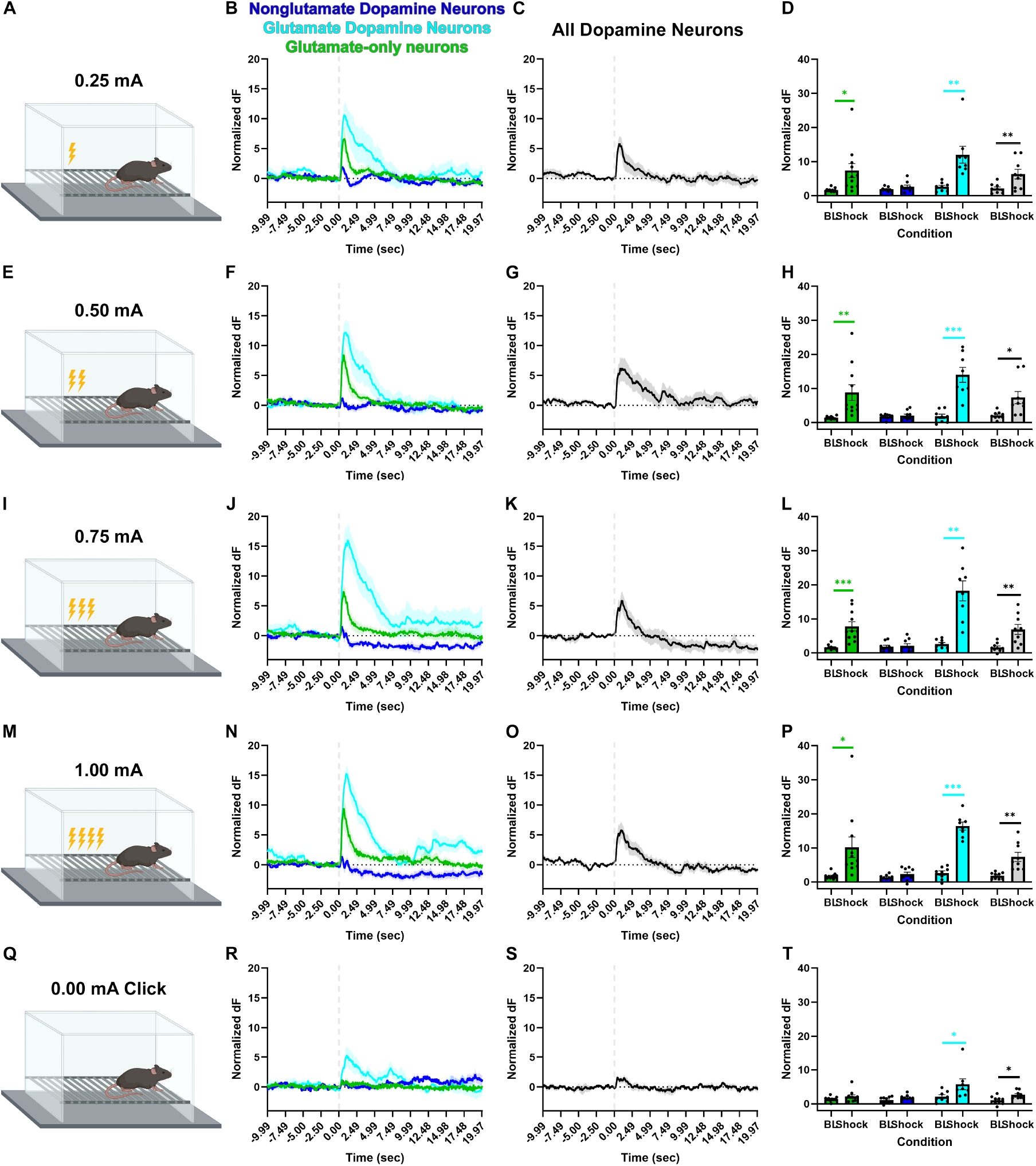
Cell-type specific aversion signaling. Neuronal activity during unsignaled footshock trials. Nonglutamate-dopamine (dark blue), glutamate-dopamine (cyan), glutamate-only (green), and all dopamine (black) neuronal activity for 0.25 mA (**A-D**), 0.50 mA (**E-H**), 0.75 mA (**I-L**), 1.00 mA (**M-P**), and 0.00 mA that caused an audible click (**Q-T**). Glutamate-only neurons: current x time interaction *F*(4,32) = 7.06, *p* = 0.019; Sidak-adjusted pairwise comparisons BL x 0.25 mA *p* = 0.018, BL x 0.50 mA *p* = 0.008, BL x 0.75 mA *p* < 0.001, BL x 1.00 mA *p* = 0.015, BL x 0.00 mA *p* = 0.104). Nonglutamate-dopamine neurons: No main effects or interactions: main effect time *F*(1,8) = 4.16, *p* = 0.760; main effect current *F*(4,32) = 0.73, *p* = 0.576; interaction *F*(4,32) = 0.61, *p* = 0.655. Glutamate-dopamine neurons current x time interaction: *F*(4,28) = 6.34, *p* < 0.001; Sidak-adjusted pairwise comparisons BL x 0.25 mA *p* = 0.004, BL x 0.50 mA *p* < 0.001, BL x 0.75 mA *p* = 0.001, BL x 1.00 mA *p* < 0.001, BL x 0.00 mA *p* = 0.043). All dopamine neurons main effect of time *F*(1,8) = 20.31, *p* = 0.002, main effect of current *F*(4,32) = 4.10, *p* = 0.009, interaction *F*(4,32) = 2.57, *p* = 0.056. BL x 0.25 mA *p* = 0.006, BL x 0.50 mA *p* = 0.015, BL x 0.75 mA *p* = 0.007, BL x 1.00 mA *p* = 0.005, BL x 0.00 mA *p* = 0.018. * *p* < 0.05, ** *p* < 0.01, *** *p* < 0.001

**Supplementary Figure 7.**
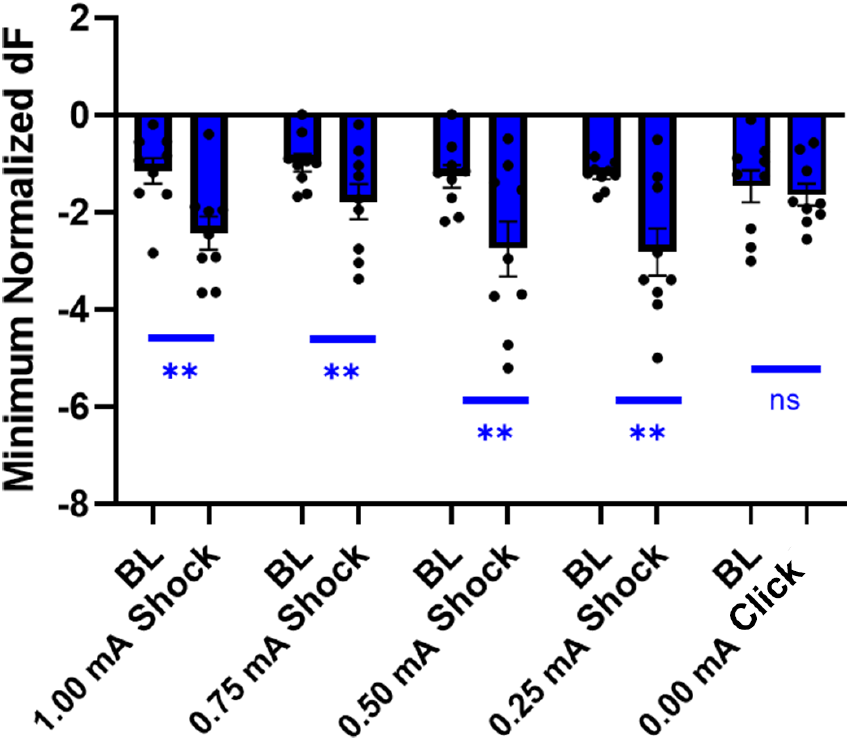
Nonglutamate-dopamine neurons reduce neuronal activity in response to footshock. Minimum normalized GCaMP response at baseline and shock delivery. Current x time interaction: *F*(4,32) = 3.23, *p* = 0.025. Sidak-adjusted pairwise comparisons BL x 0.25 mA *p* < 0.001, BL x 0.50 mA *p* = 0.040, BL x 0.75 mA *p* = 0.013, BL x 1.00 mA *p* = 0.010. ns = not significant, ** *p* < 0.01

**Supplementary Figure 8.**
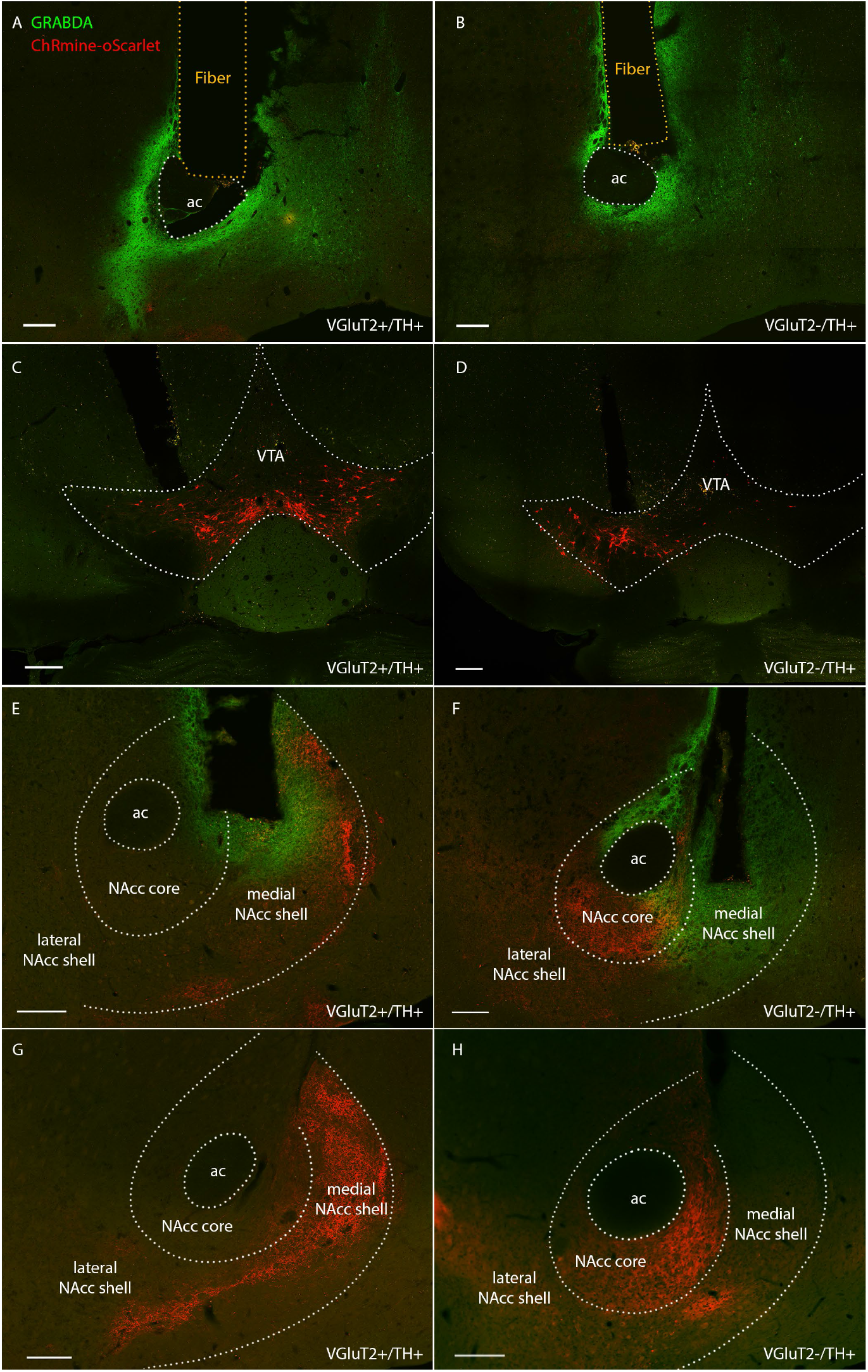
Histology for GRAB_DA_ experiments. Example optic recording fiber placement and pan-neuronal viral GRAB_DA1h_-EGFP expression in NAcc and viral ChRmine-oScarlet expression in VTA with (**A, C, E, G**) Con/Fon-ChRmine in VGluT2::Cre/TH::Flp mice targeting glutamate-dopamine neurons and (**B, D, F, H**) Coff/Fon-ChRmine in VGluT2::Cre/TH::Flp mice targeting nonglutamate-dopamine neurons. (**E, G**) Glutamate-dopamine ChRmine-oScarlet terminals are localized predominantly in NAcc shell with some terminals localized in medial NAcc core. (**F, H**) Nonglutamate-dopamine ChRmine-oScarlet fibers are localized predominantly in NAcc core with some terminals in ventral and lateral NAcc shell. All scale bars = 200 µm. ac = anterior commissure, NAcc = nucleus accumbens, VTA = ventral tegmental area.

**Supplementary Figure 9.**
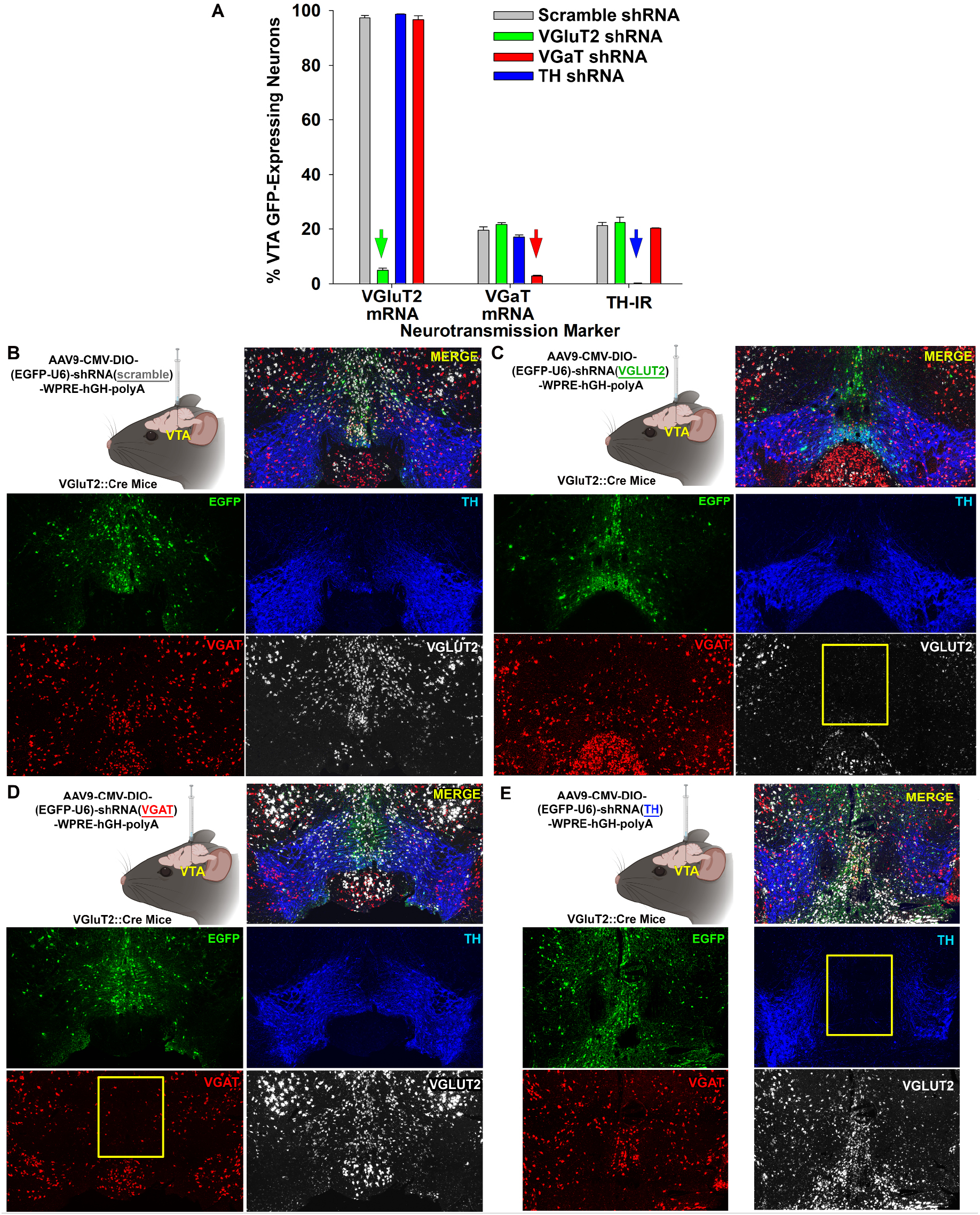
Cell-type specific neurotransmission-related knockdowns. Cre-dependent shRNA vectors were injected into VTA of VGluT2::Cre mice. VTA brain sections were labeled with *in situ* hybridization for VGluT2 and VGaT mRNA transcripts and with immunohistochemistry for TH immunoreactivity (IR). (**A**) shRNA vectors selectively and robustly silence target gene transcription while scramble control vectors have no effect. (**B-E**) Injection schema and example histology showing effectiveness of shRNA-EGFP vectors. (**B**) Scramble control vector does not reduce TH-IR, VGaT mRNA, or VGluT2 mRNA. (**C**) VGluT2 shRNA vector reduces VGluT2 mRNA, but not TH-IR or VGaT mRNA. (**D**) VGaT shRNA vector reduces VGaT mRNA, but not TH-IR or VGluT2 mRNA. (**E**) TH shRNA vector reduces TH-IR, but not VGaT or VGluT2 mRNA.

**Supplementary Figure 10.**
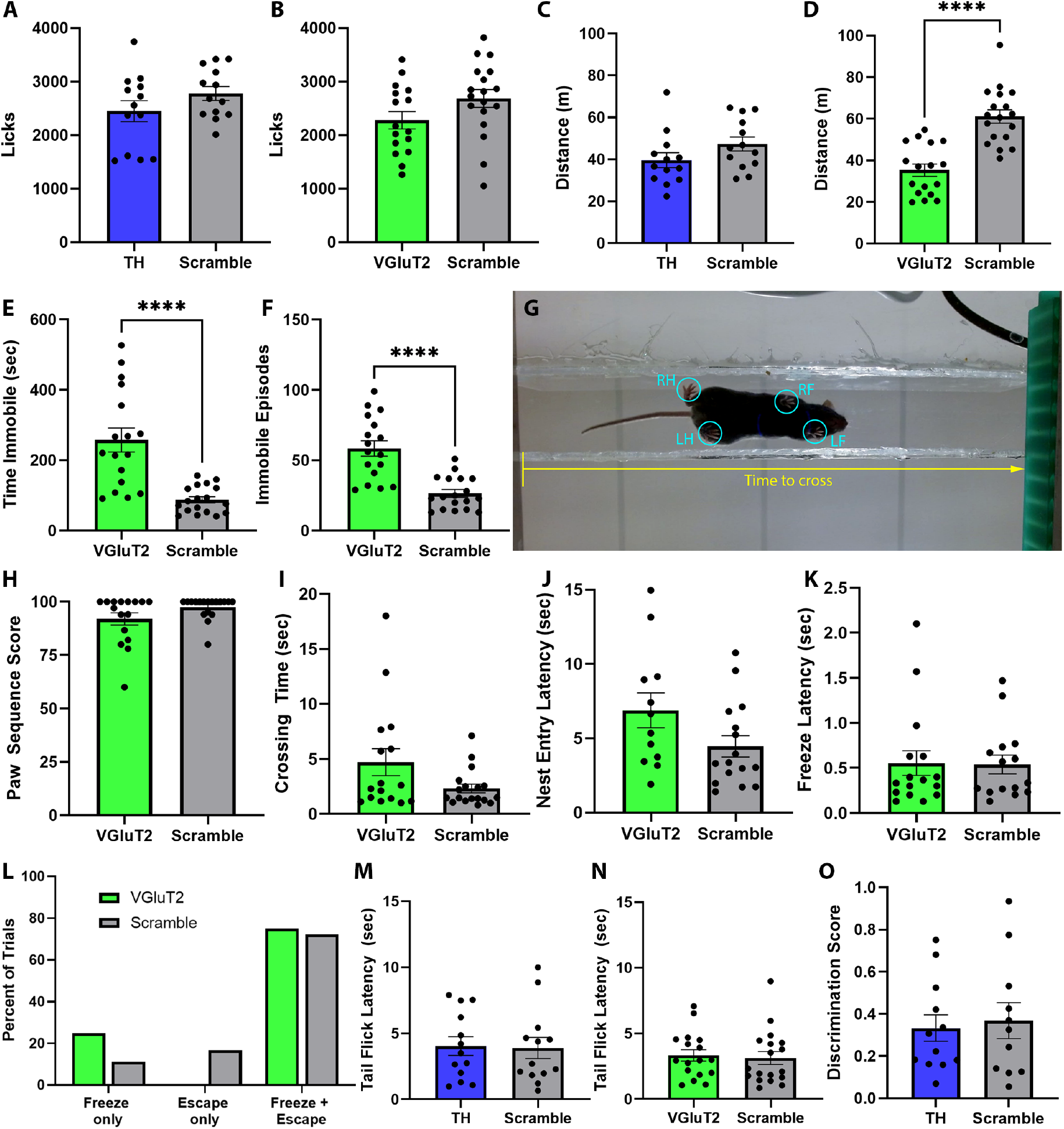
Baseline behavior analysis for shRNA knockdown experiments. Relates to Figures 6 and 7. Mice with TH knockdown in VTA glutamate neurons (TH) and mice with VGluT2 knockdown in VTA dopamine neurons (VGluT2) were compared to controls (scramble) in a battery of behavioral assays to contextualize results of conditioned reward and aversion experiments. (**A-B**) Free sucrose consumption test. (**A**) No effect of TH knockdown on number of licks, *t*(24) = 1.40, *p* = 0.174. (**B**) No effect of VGluT2 knockdown on number of licks, *t*(32) = 1.75, *p* = 0.089. (**C-F**) Open field locomotor test. (**C**) No effect of TH knockdown on distance traveled, *t*(24) = 1.59, *p* = 0.125. (**D**) Reduced distance traveled with VGluT2 knockdown, *t*(33) = 5.93, *p* < 0.0001. (**E**) Increased time spent immobile with VGluT2 knockdown, *t*(33) = 4.96, *p* < 0.0001. (**F**) Greater number of immobile episodes with VGluT2 knockdown, *t*(33) = 5.34, *p* < 0.0001. (**G-I**) Catwalk channel motor assessment. (**G**) Video frame showing catwalk channel apparatus. Mice were scored for regularity index of paw placement sequences (cyan) as well as latency to cross the catwalk (yellow). LF = left front paw, LH = left hind paw, RF = right front paw, RH = right hind paw. (**H**) No effect of VGluT2 knockdown on paw sequence regularity index score, *U* = 105.50, *p* = 0.133. (**I**) No effect of VGluT2 knockdown on latency to cross the catwalk, *U* = 106.00, *p* = 0.195. (**J-L**) Looming disc test. (**J**) No effect of VGluT2 knockdown on latency to enter the nest following loom stimulus, *U* = 57.00, *p* = 0.073. (**K**) No effect of VGluT2 knockdown on latency to freeze following loom stimulus, *U* = 112.00, *p* = 0.763. (**L**) No effect of VGluT2 knockdown on distribution of trials resulting in freezing only, escaping only (nest entry), or freezing then escaping following loom stimulus, *p* = 0.241. (**M-N**) Hot water tail immersion test. (**M**) No effect of TH knockdown on latency to remove and flick tail, *t*(24) = 0.13, *p* = 0.895. (**N**) No effect of VGluT2 knockdown on latency to remove and flick tail, *U* = 132.00, *p* = 0.503. (**O**) Novel object recognition test. No effect of TH knockdown on object discrimination score, *t*(21) = 0.33, *p* = 0.744. **** *p* < 0.0001

**Supplementary Table 1.** Electrophysiological Properties. For all tests, α was set to 0.05. Degrees of freedom (*df*) are listed as [*df* between groups, *df* within groups], where applicable. Significant *p*-values are in bold typeface. Data represent mean ± SEM. *N* for each category is in parentheses.

| **Supplementary Table 1: Electrophysiological properties** |  |  |  |  |  |
| --- | --- | --- | --- | --- | --- |
| **Property** | **Glutamate** | **Glu-DA** | **Non-Glu DA** | **ANOVA** | ***p*-Value** |
| *Capacitance (pF)* | 35.33 ± 1.387 (27) | 36.64 ± 1.341 (36) | 39.41 ± 1.851 (29) | $F(2, 89) = 1.719$ | 0.1852 |
| *Input Resistance (MΩ)* | 412.7 ± 45.96 (21) | 548.0 ± 48.14 (35) | 444.9 ± 47.78 (27) | $F(2, 80) = 2.213$ | 0.1161 |
| *Membrane Potential (mV)* | -45.0 ± 1.909 (22) | -39.35 ± 1.308 (35) | -40.15 ± 1.312 (28) | $F(2, 82) = 3.814$ | **0.0261** |

**Supplementary Table 2.** Main neural population variables and parameters. Descriptions of variables and parameters in the reinforcement learning model. Relates to Figure 8.

| **Supplementary Table 2: Main neural population variables and parameters** |  |
| --- | --- |
| $I_t$ | Input from prefrontal cortex communicating conditioned auditory stimulus |
| $u_t^d$ | GCaMP-imaged dopaminergic neural activity in ventral tegmental area (VTA) |
| $u_t^{gd}$ | Activity of co-expressing glutamate/dopaminergic VTA neurons |
| $s_t^{d1}$ | Dopamine released from pure dopamine neurons |
| $s_t^{d2}$ | Dopamine released from co-expressing neurons |
| $s_t^{dtot}$ | Total weighted sum of all released dopamine driving action-selection in striatum |
| $s_t^g$ | Glutamate released from co-expressing neurons |
| $a_t$ | Action-selection variable in striatum driven by VTA dopamine and glutamate |
| $r_t$ | Reward or shock response conditioned on action and context on each trial |
| $w_t^{di}$ | Synaptic weight of conditioned stimulus input to pure dopaminergic neural population |
| $w_t^{gdi}$ | Synaptic weight of stimulus to co-expressing dopamine-glutamate neural population |
| $w_t^{df}$ | Synaptic weight of feedback to pure dopaminergic neural population |
| $w_t^{gdf}$ | Synaptic weight of feedback to co-expressing dopamine-glutamate neural population |
| $w_t^d$ | Synaptic weight of dopamine from VTA to drive striatal action-selecting populations |
| $w_t^g$ | Synaptic weight of glutamate from VTA to drive striatal action-selecting populations |
| $\theta_t$ | Action-selection threshold, triggers nose poke or freezing when crossed |

